# B cell tolerance is dysregulated in people with complex regional pain syndrome: A high-dimensional immunophenotyping study

**DOI:** 10.64898/2026.11.18.760656

**Authors:** Ava C. Mason, Frederick A. Collin, Jackson F. Karrasch, Benjamin Heng, Seong Beom Anh, Shivani Krishnamurthy, Jayden A. O’Brien, Yun Huang, Rebecca Wong, Vaughan G. Macefield, Marc A. Russo, Nick Christelis, Peter Georgius, Joanne H. Reed, Paul J. Austin

**Author notes:** Corresponding author: Paul J. Austin, Brain and Mind Centre, 100 Mallet St, Camperdown, NSW, 2050, Australia.

## Abstract

Complex regional pain syndrome (CRPS) is a devastating chronic pain disorder that typically arises after distal limb trauma. It has a complex, multi-system pathophysiology that is not well understood, preventing the development of targeted therapies. Observations of self-reactive antibodies in CRPS patients that can elicit pain hypersensitivity in rodents have established an autoimmune contribution to the onset and maintenance of CRPS-like pain, although the mechanisms underlying this remain unclear. To understand the development of pathogenic autoantibody responses in CRPS, this study performed the first high-dimensional characterisation of B cell activity in a cohort of 42 CRPS patients and 42 matched healthy controls. Immunophenotyping of peripheral immune cell populations by mass cytometry identified altered frequencies of class-switched and double-negative memory B cell subsets in CRPS, and increased use of autoimmune-associated IGHV1-69 and IGHV4-34 gene-encoded heavy-chains within the class-switched B cell compartment. Additional proteomic and autoantibody assessments identified differences in serum protein and IgA autoantibody profiles that may lead to altered complement signalling and pro-inflammatory cytokine production in CRPS. Together, these findings represent evidence of dysregulated B cell self-tolerance in CRPS that could facilitate production of pathogenic autoantibodies in people with the condition.

## Introduction

Complex regional pain syndrome (CRPS) is a severe and highly debilitating chronic pain disorder that usually affects a distal limb after tissue trauma, such as fracture, sprain, or surgery. Primarily characterised by unresolving regional pain that is disproportionate to the inciting injury, CRPS is considered one of the most painful conditions humans can experience [1, 2]. It is further associated with various localised sensory and non-sensory features including hyperalgesia (increased sensitivity to painful stimuli) and allodynia (sensitivity to non-painful stimuli), altered skin temperature and colour, increased sweating and swelling, abnormal nail or hair growth, motor disturbances, and decreased range of movement [3]. Many patients experience significant functional impairment and reduced quality of life as a result, along with anxiety, depression, and even suicidal ideation [4, 5, 1]. Existing treatment strategies are ineffective at relieving this burden, with a recent Cochrane review finding no high-certainty evidence that any of the current interventions for CRPS reliably reduce pain or disability [6].

The development of mechanism-based therapeutics for CRPS is prevented by our incomplete understanding of its complex pathophysiology, which involves both neurological and inflammatory dysfunction. CRPS is now also believed to have a substantial autoimmune component, based on observations of autoreactive IgG and IgM in patients that can produce a CRPS phenotype when passively transferred to rodents with a hindlimb injury [7–12], which constitutes indirect evidence of an autoimmune aetiology according to current criteria [13]. This has prompted further enquiry into autoantibody-mediated mechanisms of CRPS pain, mostly involving *in vitro* profiling of autoantibodies against specific tissue targets, which has revealed that subgroups of CRPS patients display levels of autoantibodies against autonomic receptors on neurons and other native antigens [14–20], although the clinical relevance of these targets remains unknown [21]. Further studies have identified persistent changes to the broader immune landscape in CRPS that could facilitate pathogenic autoimmune responses, including pro- and anti-inflammatory cytokine dysregulation [22, 23] and altered T cell ratios [24–27].

Surprisingly, peripheral B cell populations are yet to be profiled in clinical CRPS, despite the well-established role of autoantibodies in the condition. Preclinical models of CRPS and neuropathic pain support a B cell involvement in CRPS-like symptoms, demonstrating that post-injury pain hypersensitivity in rodents is not only driven by immunoglobulin deposition at central and peripheral nervous system sites, but can be alleviated or completely prevented by inhibiting B cell activity via anti-CD20 treatment or genetic depletion, suggesting a direct B cell contribution to CRPS-like pain [11, 28–30].

In light of these observations, we hypothesise that people with CRPS will have discernible differences in their circulating B cell repertoires that could facilitate pain-promoting autoantibody responses. To characterise these changes, we used single-cell resolution suspension mass cytometry to profile major circulating B cell subsets in a cohort of 42 CRPS subjects and 42 matched healthy donors. Serum proteomic analysis and a high-scope autoantibody array were also used for further exploration of the peripheral immune environment in CRPS. This represents the first comprehensive assessment of B cell subset differentiation and maturation and associated peripheral immune changes in clinical CRPS.

## Methods

### Participants and sample collection

The study included 42 individuals with CRPS (31 women), and 42 pain-free healthy donors who were matched to patients by age and sex. Exclusion criteria were as follows: under the age of 18; pregnant; less than 3 months post-onset (CRPS group); or experiencing any acute pain (healthy control group). All CRPS subjects met the Budapest criteria for diagnosis [31], and all participants provided written informed consent. Samples were collected by trained phlebotomists at Neuroscience Research Australia (Sydney, NSW), Genesis Research Services (Broadmeadow, NSW), or the Baker Heart and Diabetes Institute (Melbourne, VIC). For each participant, 10 mL whole blood was collected into an EDTA tube via venepuncture to the antecubital fossa (unaffected limb for CRPS subjects). Tubes were immediately mixed by gentle inversion to prevent coagulation, then 0.5 mL aliquots were transferred to 1.5 mL cryovials containing 0.7 mL PROT1 stabilisation buffer (SmartTube Inc.) and mixed by inversion. Samples were incubated at RT for 10 minutes before being transferred to -80°C for long-term storage. An additional 10 mL of whole blood was collected into a serum tube and allowed to coagulate before centrifugation and collection of the serum supernatant into 1.5 mL cryovials, which were also then transferred to -80°C for long-term storage. Participants were asked to cease any immunomodulating medications (NSAIDs, corticosteroids, opioids) for at least 7 days prior to sample collection (although three CRPS participants reported NSAID use during this period). While a longer or stricter drug washout would have been ideal for limiting residual effects on immune cell populations, this was not possible due to practical and ethical considerations.

### Pain and psychological profiling

To assess pain and psychological symptoms, CRPS participants rated their pain severity at the time of sample collection on a Visual-Analogue Scale (VAS), and also completed the following questionnaires: revised Short-Form McGill Pain Questionnaire (SF-MPQ-2) [32], Depression, Anxiety, and Stress Scale (DASS) [33], Pain Catastrophising Scale (PCS) [34], Tampa Scale for Kinesiophobia (TSK) [35], Pain Self-Efficacy Questionnaire (PSEQ) [36]. Participants in the Melbourne cohort also completed the CRPS Symptoms Scale (CSS) [37]. Final scores were interpreted according to published guidelines [36, 38–40].

### Mass cytometry

#### Staining and acquisition

Peripheral blood samples from CRPS patients and controls were analysed by two mass cytometry studies. The preliminary study profiled B cells and T cells with a panel of 39 metal-conjugated antibodies (Supplementary Table 1), and included samples from the 27 CRPS participants (*n* = 26 acquired) collected between 2022 and 2024 at the Baker Heart and Diabetes Institute (Melbourne, NSW). The same samples were profiled in the follow-up study with an extended B cell panel (Supplementary Table 2), along with an additional 15 CRPS samples collected between 2017 and 2018 at Genesis Research Services (Broadmeadow, NSW). For both experiments, antibody concentrations were optimised through multiple test titrations performed on healthy donor blood samples (Supplementary Figure 1). Experimental samples were prepared, stained, and acquired in multiple batches according to the protocol outlined in Supplementary Table 3, which was adapted from our previous studies [41, 42, 24]. All samples were acquired by cytometry by time-of-flight (CyTOF) on a Helios suspension mass cytometer (Fluidigm). The full staining and acquisition protocol is provided in Supplementary Methods 1.

#### Manual gating analysis

Cytometry data were first analysed by manual gating in FlowJo (v10.10.1, BD) according to the strategy outlined in Supplementary Figure 2. Bead-normalised .fcs files were imported into FlowJo and concatenated where necessary, then pre-gated to remove doublets, de-barcode paired samples, remove non-leukocytes and neutrophils, and distinguish B cell and T cell populations. Major B cell and T cell subsets were then gated according to core surface marker expression (Supplementary Figure 2; Supplementary Table 4), and cell statistics for each gated subset were exported for statistical analysis.

#### Unsupervised analysis

Pre-gated CD19^+^ and CD3^+^ populations were exported from FlowJo as .fcs files and analysed using the TidyTOF *R* package [43] in RStudio. First, raw single-cell data were arcsinh-transformed with a cofactor of 5 (the standard method for CyTOF data [44]), then batch-corrected via linear rescaling. FlowSOM clustering and dimensionality reduction were then performed according to the parameters outlined in Supplementary Table 5. Heatmaps and ridgeline plots were used to identify clusters based on core surface marker expression (Supplementary Tables 4-5). Duplicate clusters were merged, and clusters containing doublets or heterogeneous populations were removed. Differential abundance and expression analyses were then performed on the merged metaclusters.

### Proteomics

Serum samples from CRPS participants (*n* = 42) and healthy controls (*n* = 20) were assessed by SWATH mass spectrometry (MS)-based proteomic analysis, conducted according to the parameters outlined in Supplementary Methods 2. Proteins with a fold change (FC) > ±1.5 and P < 0.05 for CRPS compared to controls were considered differentially expressed [45, 46]. To understand the functional significance of the differentially expressed proteins, gene set enrichment analysis (GSEA) was performed using the Gene Ontology (GO) biological process and Jensen DISEASES databases [47, 48], and pathway analysis was conducted using the Reactome Pathways library [49]. All downstream analyses of proteomic findings were performed using EnrichR (https://maayanlab.cloud/Enrichr) [50].

### Autoantibody profiling

Serum IgG and IgA autoantibodies from CRPS subjects (*n* = 42) and healthy controls (*n* = 18) were profiled using the iOme^®^ Discovery Protein Array, a high-density, slide-based protein microarray that uses KREX^TM^ protein folding technology to profile serum autoantibodies against over 1800 correctly folded, full-length human proteins, including established and putative autoantigens (https://somalogic.com/krex-protein-list/). The assay and corresponding data analysis was performed by Houston Omics Collaborative at the University of Houston (Houston, USA) according to previously described methods [51]. Data were analysed by penetrance fold change (pFC) testing, which involves generating a per-antigen baseline against the reference group (e.g. healthy donors for CRPS vs. control) using normalised and filtered input data, then calculating the fold-change (FC) for each antigen in each sample relative to the baseline value. Group comparisons were then performed on the per-antigen frequency (penetrance frequency) of the samples with FC ≥ 2.0 relative to the per-antigen baseline value. Fisher’s exact test was then used to compare groups with P < 0.2.

### Statistical analysis

Statistical analysis and data visualisation were performed in R (v.4.5.3) or Prism 11 (GraphPad Software Inc.). Cohort age and sex statistics were compared using Welch’s t-tests and Fisher’s exact tests, respectively. Differential cell abundance comparisons were performed using Welch’s t-tests for parametric data (determined with Shapiro-Wilk tests) or Mann-Whitney U-tests for non-parametric data. Differential expression analysis was conducted using the TidyTOF R package’s implementation of diffcyt’s *limma* method to = 0.05 [52]. Associations between cell subset abundance and clinical variables (age, sex, CRPS duration, affected limb, pain and psychological symptoms) were assessed with simple linear regression analysis and Pearson correlation analysis, using the Benjamini-Hochberg false discovery rate method used for multiple comparisons correction. Statistical significance was reported for mass cytometry data when P < 0.05 for manual gating analysis, or multiple comparisons-corrected P_adj_ < 0.05 for all other analyses.

### Study approval

This study was approved by the University of Sydney Human Research Ethics Committee (HREC #2021/083). All participants provided written informed consent.

## Results

### Participant demographics and clinical findings

The CRPS cohort included 31 women and 11 men, and had an average age of 45.8 years. This reflects the typical age at onset and the 2-3 times higher female prevalence of CRPS reported by large population studies [53, 54]. Pain-free healthy donors were matched to CRPS patients by sex and age, resulting in the same proportion of female to male participants and a mean age of 43.3 years (P = 0.472). Smaller control cohorts were used for the proteomic and autoantibody screens, and three subjects (two controls, one CRPS) were excluded from mass cytometry analysis due to poor sample quality, leading to slight variations in cohort age and sex ratios, but these were statistically insignificant across all studies (Supplementary Table 6).

Within the CRPS cohort, the most common inciting injury was distal limb fracture (*n* = 16), followed by surgery (*n* = 8) or soft tissue injury (*n* = 8). Time since onset ranged from 4 months to 24.4 years, with an average of 5.1 years. Nearly three-quarters of the cohort (73%) had ‘persistent’ CRPS, where duration exceeded 18 months. These findings are summarised in Table 1, with extended clinical profiles of each CRPS participant provided in Supplementary Table 7. Results of pain and psychological symptoms assessment in the CRPS cohort are also outlined in Table 1. Participants reported a moderate level of pain on the day of sample collection, with an average VAS score of 4.8/10. Moderate pain intensity was confirmed by the SF-MPQ-2, where the mean score was 4.5/10. However, the Melbourne cohort scored 6.1/8 on the CSS (patient-reported section), indicating severe CRPS. The complete CRPS cohort also experienced high levels of pain catastrophising based on the PCS, and a severe fear of movement according to the TSK, with only a moderate level of pain self-efficacy based on the PSEQ. Patient DASS scores indicated mild levels of depression and stress, along with moderate anxiety. Collectively, these findings demonstrate the clear impact of ongoing pain on the mental and emotional wellbeing of individuals with CRPS. Notably, some psychological measures differed between female and male CRPS participants (Figure 1). On average, women with CRPS reported significantly lower DASS stress levels than men (P = 0.0183; Fig. 1C), scoring in the normal range for this subscale, while male patients scored in the moderate range. This disparity did not appear to reflect differences in pain intensity or CRPS symptoms, as both groups scored similarly on the VAS, SF-MPQ-2, and CSS (P = 0.6693; P = 0.8613; P = 0.0937; Fig. 1G-I). Together, these findings highlight the role of sex and gender in the perception of and response to ongoing pain, which may underpin differences in how men and women with CRPS experience or describe the psychological impact of their pain.

**Figure 1.**
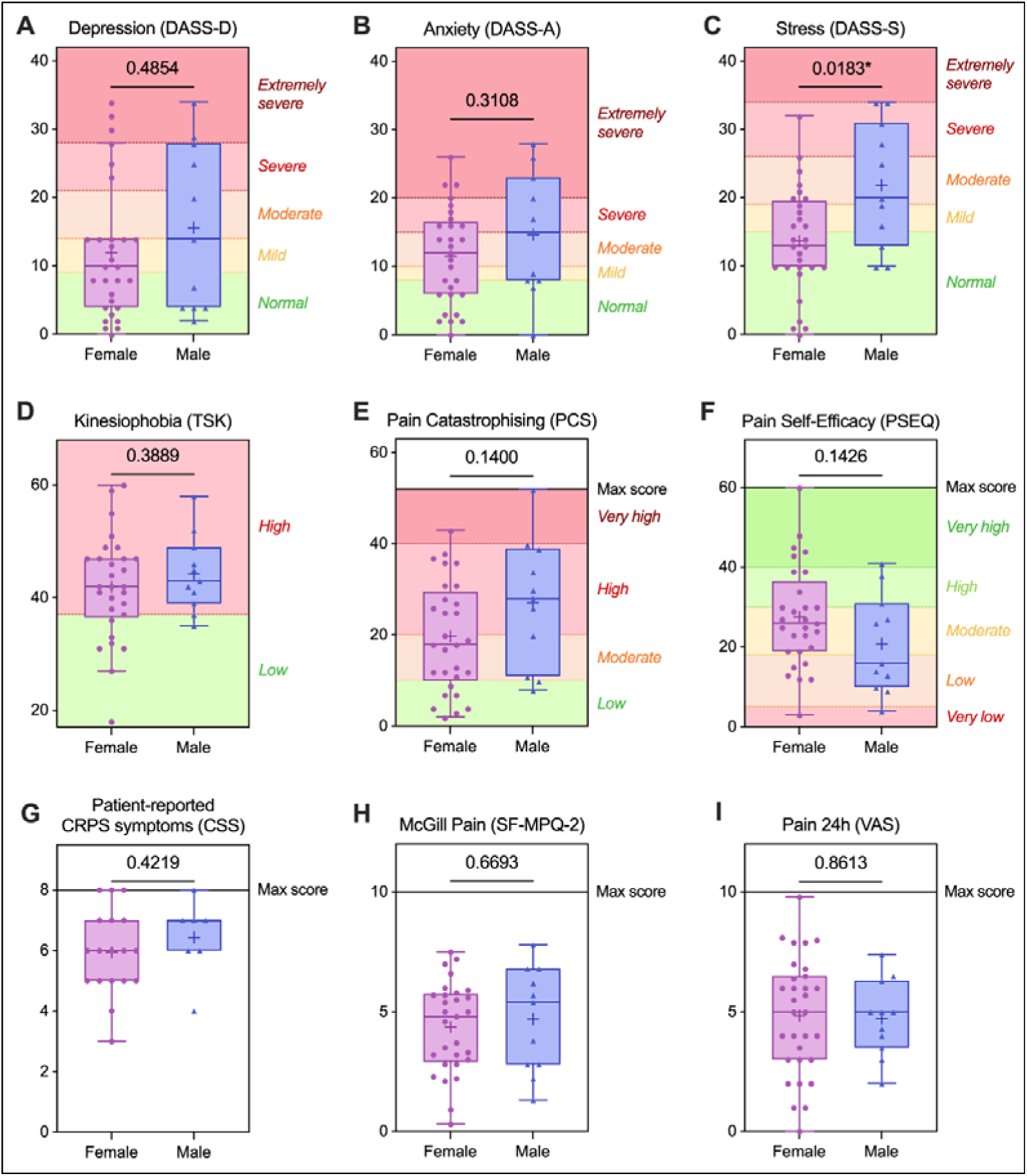
Sex differences in psychological symptoms reported by CRPS participants. Women with CRPS (*n* = 31) had lower DASS stress scores than men with CRPS (*n* = 10; **C**), despite reporting similar pain intensity and CRPS symptoms (**G**-**I**). Boxplots show interquartile range (IQR), median (solid lines), and mean (+) for each scale, with whiskers extending to 1.5x IQR. Each point represents an individual CRPS participant (*n* = 42 for VAS pain, *n* = 41 for all other scales). Solid y-axis lines in panels **E**-**I** indicate maximum possible score for a given scale; coloured segments divided by dashed lines in panels **A-F** represent thresholds for final score interpretation, according to published guidelines for each scale [36, 38–40]. Statistical comparisons were performed using Welch’s t-tests for parametric data (determined by Shapiro-Wilk test) or Mann-Whitney U-tests for non-parametric data, with *P < 0.05. CSS = CRPS Symptoms Scale; DASS = Depression, Anxiety, Stress Scale; PCS = Pain Catastrophising Scale; PSEQ = Pain Self-Efficacy Questionnaire; SF-MPQ-2 = revised Short-Form McGill Pain Questionnaire; TSK = Tampa Scale for Kinesiophobia; VAS = Visual-Analogue Scale.

**Table 1.**
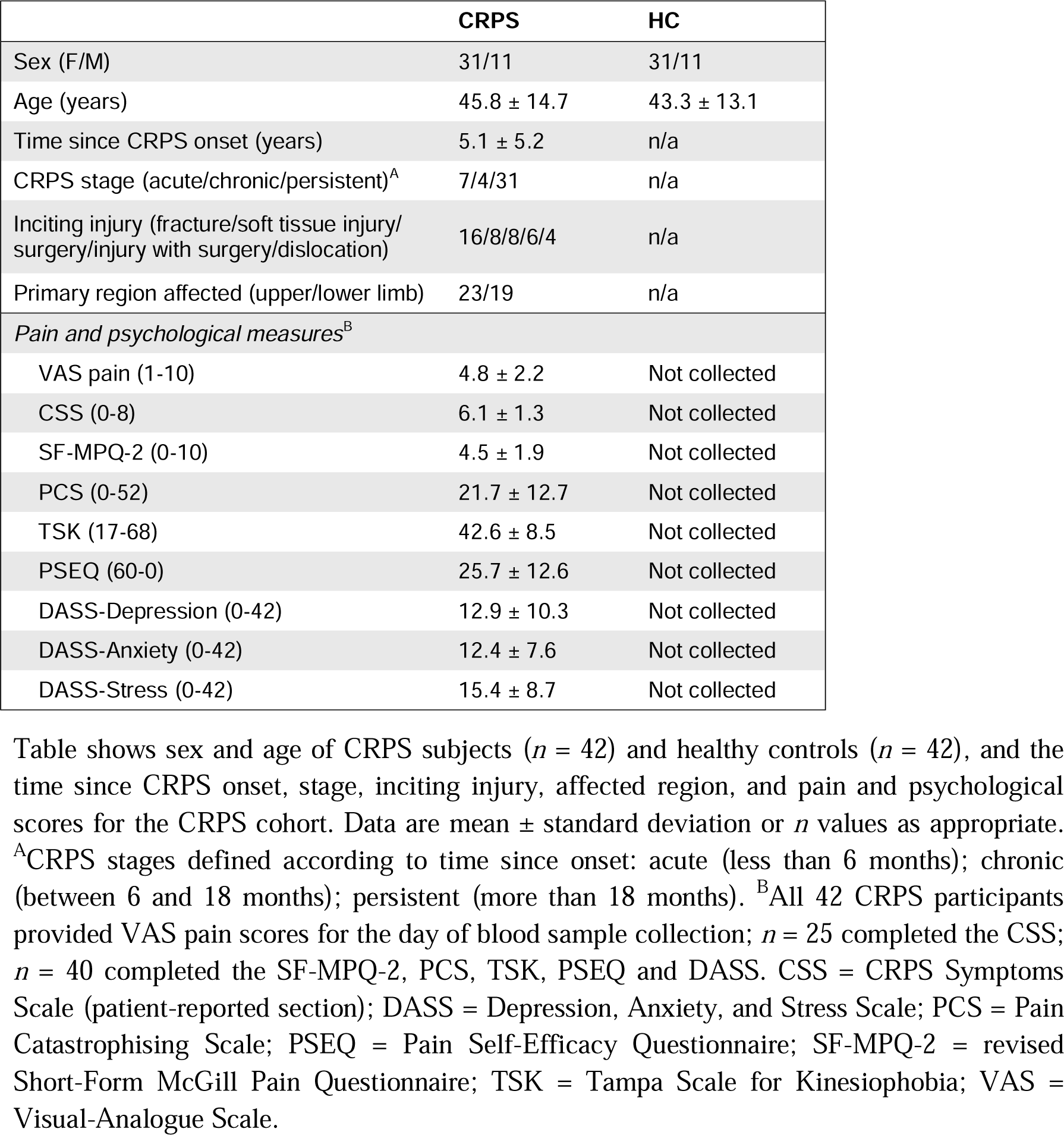
Demographics and clinical features of the CRPS and healthy control cohorts.

### CRPS participants showed changes in memory B cell subset frequencies and expression of B cell regulation and tolerance markers

Circulating B cell subsets were analysed by both manual gating in FlowJo and unsupervised FlowSOM clustering. Manual analysis of the class-switched memory B cell (CSM) compartment revealed a proportional reduction in resting-state CSM B cells (IgD^−^ CD38^lo^CD27^+^CD21^+^) in CRPS (P = 0.0006; Figure 2A), although this change was not observed during unsupervised analysis (likely reflecting slight differences in the surface markers used to phenotype this population between analysis methods; Supplementary Figure 4B). CRPS participants also showed differences in distinct ‘double-negative’ (DN) memory B cell populations compared to healthy controls. DN2 memory B cells, defined as IgD^−^ CD27^−^CD21^−^CD38^−^ [55, 56], were expanded in CRPS, (P = 0.012; Fig. 2B), as were FCRL4^+^ DN2 B cells, a rare ‘exhausted’ atypical memory population [56–58] that comprised less than 2% of all B cells in the control group but close to 5% in the CRPS group (P = 0.0216; Fig. 2C). Four different DN memory populations were also identified by FlowSOM clustering, with varying levels of CD38 and CD21 expression. One of these populations, a DN2-like subsets (IgM^−^IgD^−^CD27^−^CD21^−^CD38^lo^), was increased in CRPS (P_adj_ = 0.0353; Supplementary Fig. 4B).

**Figure 2.**
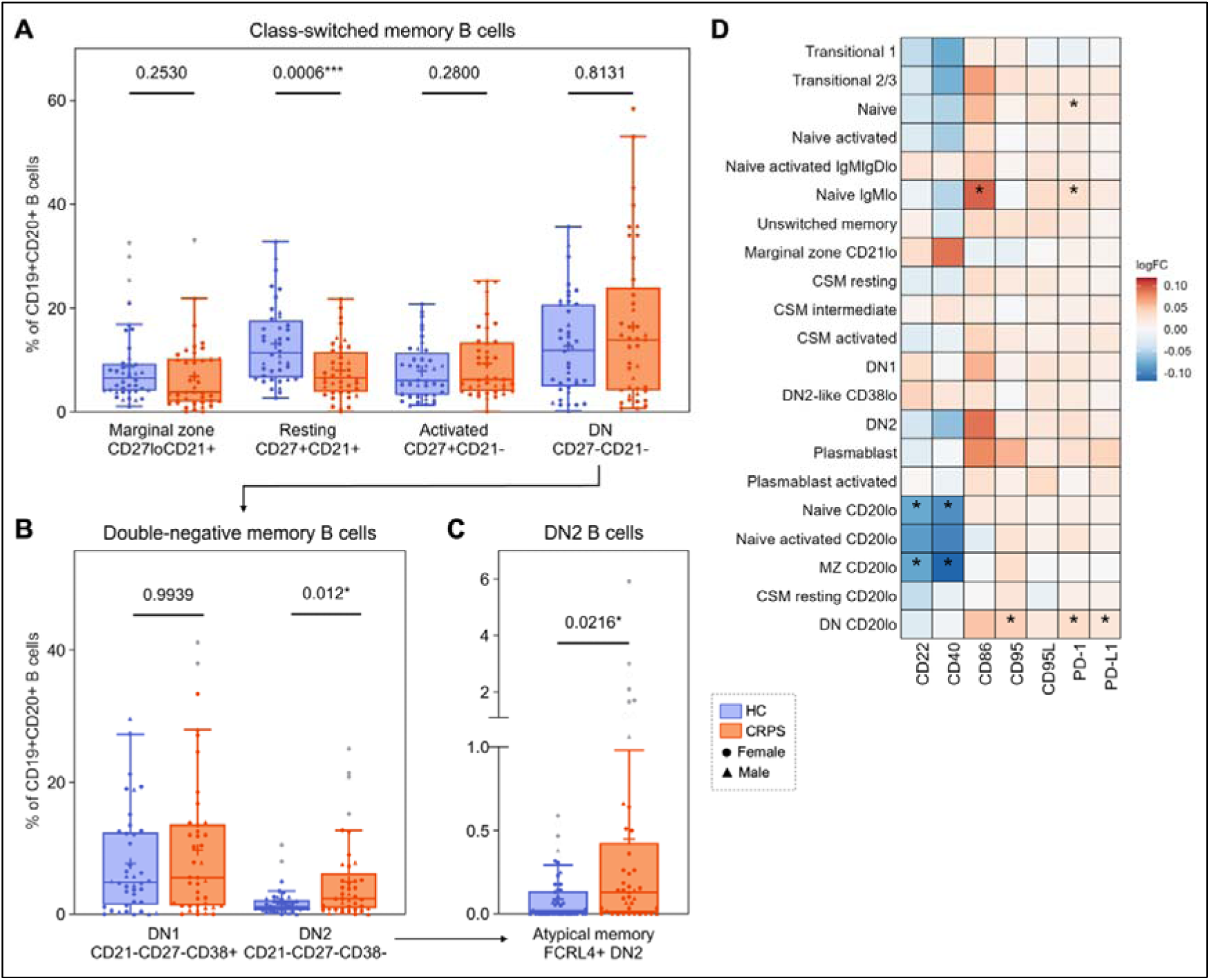
Differences in class-switched memory B cell profiles and activity marker expression between CRPS subjects (*n* = 41) and healthy donors (*n* = 40). **(A)** Manual gating analysis revealed a decreased proportion of resting memory B cells (IgD^−^CD38^−^ CD27^+^CD21^+^) in CRPS, and increased proportions of **(B)** double-negative type 2 B cells (IgD^−^CD27^−^CD21^−^CD38^−^) and **(C)** an FCRL4^+^ DN2 subset (atypical/exhausted memory). Boxplots show IQR, median (solid lines), and mean (+) for each B cell subpopulation, with whiskers extending to 1.5x IQR. Each point represents an individual participant. Arrows indicate gated parent populations passed on for subsequent analysis. Statistical comparisons were performed using Welch’s t-tests for parametric data (determined by Shapiro-Wilk test) or Mann-Whitney U-tests for non-parametric data, with *P < 0.05; ***P < 0.001. Statistical outliers (grey points) were excluded using ROUT (Q = 1%). **(D)** Distinct B cell populations also showed differential expression of regulatory markers in CRPS. Heatmap shows log-fold changes in activity marker expression for CD19^+^ FlowSOM metaclusters between CRPS and controls, with a ratio of 0 (white) indicating equal expression, a ratio greater than 0 (orange/red) indicating CRPS>control, and a ratio less than 0 (blue) indicating CRPS<control, with *P_adj_ < 0.05.

FlowSOM clustering also identified five distinct B cell populations expressing low levels of CD20 that were otherwise phenotypically identical to other B cell subsets, including a naive (IgM^+^IgD^+^CD27^−^CD21^+^CD38^lo^), activated naive (IgM^+^IgD^+^CD27^−^CD21^−^CD38^−^), marginal zone (IgM^+^IgD^−^CD27^lo^CD21^+^CD38^−^), resting CSM (IgM^lo^IgD^−^CD27^+^CD21^+^CD38^−^CD95^−^CD86^−^) and DN memory population (IgM^−^IgD^−^CD27^−^CD21^−^CD38^+^CD95^−^CD86^−^). The proportions of these CD20^lo^ subsets were unchanged between CRPS and control groups (Supplementary Fig. 4B), but they exhibited different expression profiles for multiple regulatory markers (Fig. 2D). The CD20^lo^ naive and CD20^lo^ marginal zone populations expressed lower levels of the inhibitory co-receptor CD22 (P_adj_ = 0.00713; 0.0294) and the co-stimulatory receptor CD40 (P_adj_ = 0.0294; 0.00713) in CRPS, and the CD20^lo^ DN memory subset upregulated three markers representing distinct peripheral tolerance checkpoints: PD-1 (P_adj_ = 0.00713), PD-L1 (P_adj_ = 0.0294) and CD95 (P_adj_ = 0.0410). Two other CD19^+^CD20^+^ metaclusters also showed higher PD-1 expression in CRPS, the naive population (IgM^+^IgD^+^CD27^−^CD21^+^CD38^int^; P_adj_ = 0.0447; Fig. 3D) and an IgM^lo^ anergic-like naive population (P_adj_ = 0.0294), which also upregulated the co-stimulatory marker CD86 (P_adj_ = 0.0410). The changes in CD22 expression observed during unsupervised analysis prompted further analysis of this marker using manual gating (Supplementary Figure 5), revealing a subgroup of the CRPS cohort with a significantly reduced proportion of CD22^+^ B cells compared to controls (P = 0.0276). Overall, unsupervised and manual expression analysis revealed CRPS-specific changes in B cell activity markers related to immune regulation and self-tolerance.

**Figure 3.**
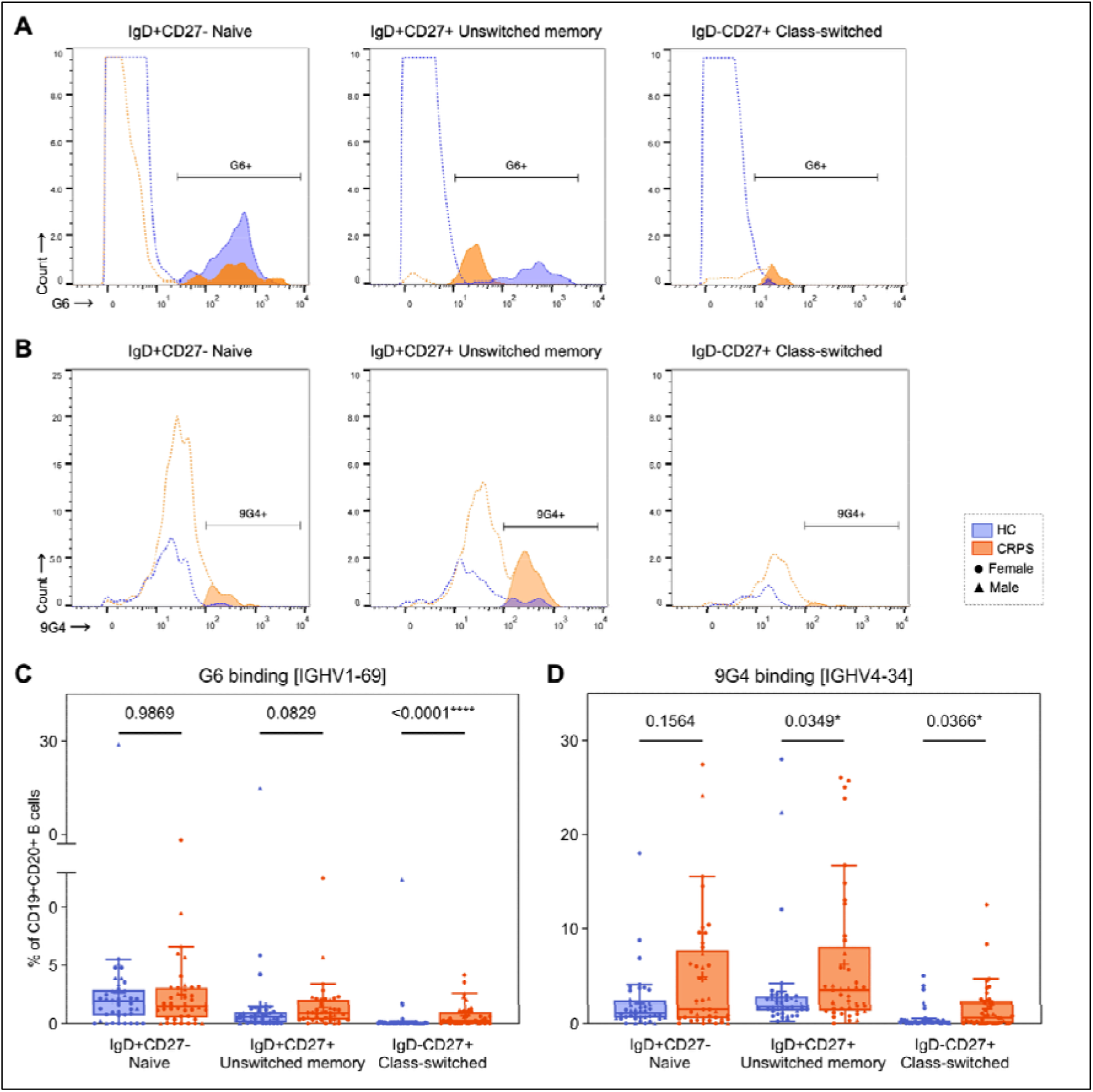
Increased G6 [IGHV1-69] and 9G4 [IGHV4-34] binding in the class-switched B cell compartment of CRPS subjects (*n* = 41) compared to healthy donors (*n* = 40). Histograms show representative manual gating strategies used to assess **(A)** G6 and **(B)** 9G4 binding across the B cell compartment between CRPS and control samples. Dotted lines represent the parent population gate in each sample; solid lines with filled areas represent its positive subpopulation gated as either G6^+^ or 9G4^+^. **(C-D)** Statistical comparisons were performed using Welch’s t-tests for parametric data (determined by Shapiro-Wilk test), or Mann-Whitney U-tests for non-parametric data, where *P < 0.05; ****P < 0.0001. Boxplots show IQR, median (solid lines), and mean (+) for each cell subpopulation, with whiskers extending to 1.5x IQR. Each point represents an individual participant.

Associations between CD19^+^ metaclusters and clinical features of the CRPS cohort (age, sex, time since onset, pain and psychological measures) were assessed using Pearson’s correlation analysis or Welch’s t-tests for categorical data, such as medication use (Supplementary Figures 7-9). The proportion of transitional type 1 B cells correlated negatively with SF-MPQ-2 scores (R = -0.613, P_adj_ = 0.0050), and VAS pain scores showed negative correlations with two DN memory B cell metaclusters: DN1 (R = -0.552, P_adj_ = 0.0068) and CD38^lo^ DN2-like (R = -0.451, P_adj_ = 0.0455), and a weak positive correlation with the CD20^lo^ naive B cell metacluster (R = 0.510, P_adj_ = 0.0098; Supplementary Fig. 7A). Multiple B cell metaclusters were proportionally reduced in CRPS participants who reported regular NSAID use compared to those who did not, including transitional 1 (P_adj_ = 0.0053), DN2 memory (P_adj_ = 0.0090), and three of the CD20^lo^ populations (Supplementary Fig. 8). In summary, the proportions of transitional, DN memory, and CD20^lo^ B cell subsets showed associations with patient-reported pain severity and varied by NSAID usage.

### Heavy-chain gene segments associated with autoimmune disease were increased in the class-switched B cell compartment in CRPS

G6 and 9G4 antibodies were used to assess expression of two heavy-chain gene segments enriched in autoimmune disease, IGHV1-69 and IGHV4-34, before and after B cell class-switching (Figure 3A). G6 and 9G4 expression in the naive B cell compartment (IgD^+^CD27^−^) was not different between CRPS and control participants (Fig. 3B-C), but the CRPS cohort showed significantly higher levels of G6 and 9G4 binding in the class-switched memory B cell compartment (IgD^-^CD27^+^) compared to controls (P < 0.0001; P = 0.0366; Fig. 2B-C), where the proportion of class-switched G6^+^ or 9G4^+^ B cells was close to zero. Unswitched or ‘innate-like’ memory B cells (USM; IgD^+^CD27^+^) also showed increased 9G4 binding in CRPS (P = 0.0349; Fig. 3C). Within the CRPS cohort, female participants displayed higher 9G4 binding in the USM and class-switched B cell compartments compared to males (P_adj_ = 0.0128; P_adj_ = 0.0364; Supplementary Figure 6). No further correlations were observed between G6 and 9G4 binding and other clinical features of the CRPS cohort (Supplementary Figure 7).

### Distinct helper T cell subsets shifted towards antigen-experienced phenotypes and showed enhanced inflammatory signalling in CRPS

Major T cell subsets were assessed during the preliminary CyTOF study by manual gating and unsupervised analysis. Naive Th cells (CD4^+^CD25^−^CD62L^+^CD45RA^+^) were proportionally reduced in CRPS compared to healthy donors (manual P = 0.0252; Figure 4A; unsupervised P_adj_ = 0.0039; Supplementary Figure 10), while effector Th cells (CD4^+^CD25^−^ CD62L^−^CD45RA^−^) were expanded (manual P = 0.0066; unsupervised P_adj_ = 0.0281). Manual gating analysis revealed that CRPS participants had an increased proportion of effector memory Th cells (CD4^+^CD25^−^CD62L^−^CD45RA^−^) compared to controls (P = 0.0176; Fig. 4A), and a reduced proportion of naive Tregs (CD4^+^CD25^+^FOXP3^+^CD62L^+^CD45RA^+^; P = 0.0369; Fig. 4B), although these populations were unchanged between groups during unsupervised analysis (Supplementary Fig. 10B). These discrepancies are likely due to slight differences in surface phenotyping for these T cell subsets between the two analysis methods (Supplementary Tables 4-5). Additional T cell populations identified by FlowSOM clustering include a CD8a^+^ Temra-like phenotype (CD8a^+^CD62L^+^CD45RA^lo^CD45RO^lo^), and a naive CD4^+^CD8a^+^ population (CD62L^+^CD27^+^CD45RA^+^CD45RO^−^), both of which were reduced in CRPS (P_adj_ = 0.0150; P_adj_ = 0.0228; Supplementary Fig. 10B). Overall, CRPS patients displayed a shift away from the naive and towards effector and effector memory subsets in the helper T cell compartment, reduced naive Tregs, and further decreases in minor T cell phenotypes that were identified by unsupervised clustering.

**Figure 4.**
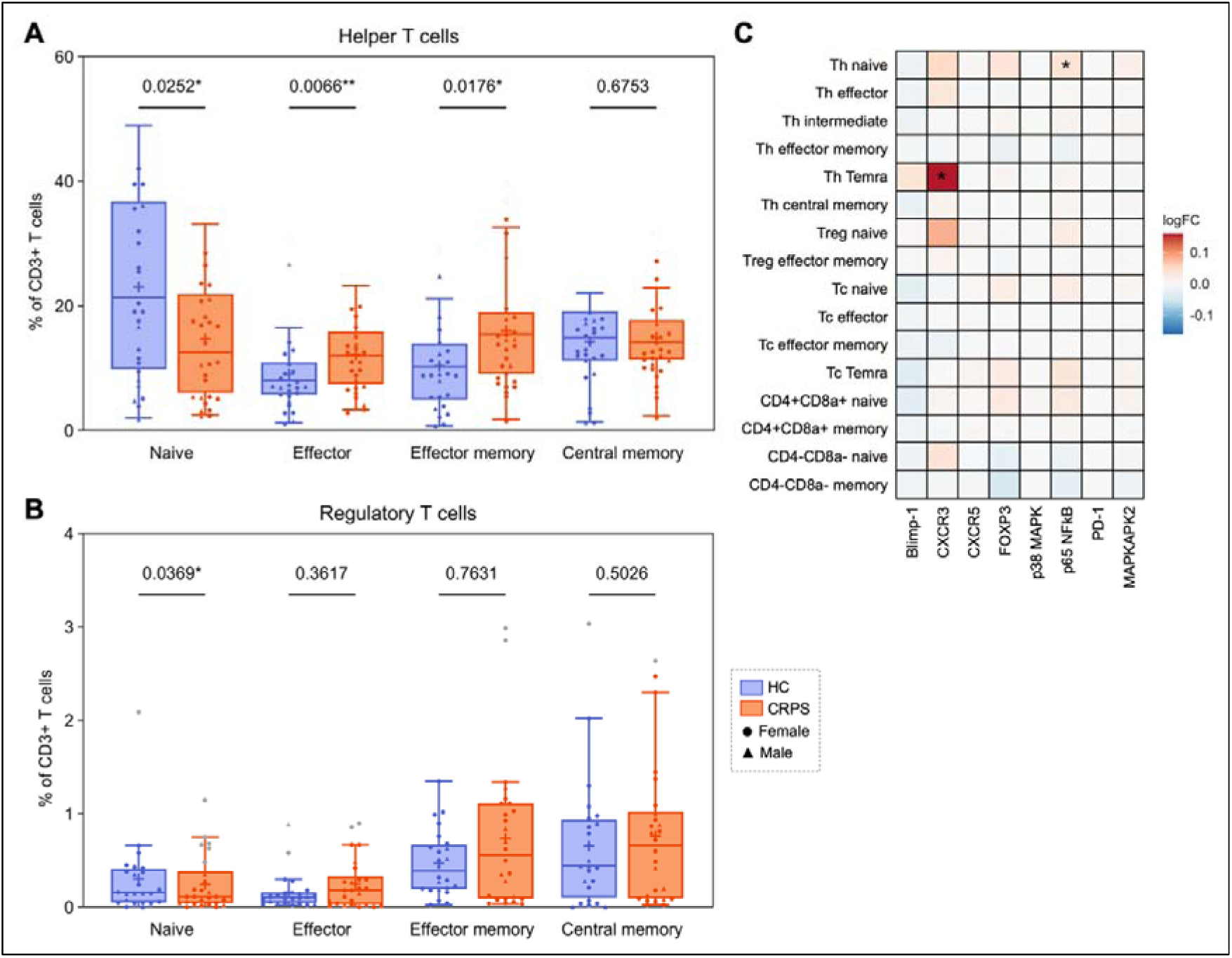
Differences in helper T cell subpopulation frequencies and inflammation marker expression between CRPS subjects (*n* = 26) and healthy donors (*n* = 24). **(A)** Manual gating analysis of the helper T cell compartment (CD4^+^CD8a^−^CD25^−^) identified a reduction in naive Th cells (CD62L^+^CD45RA^+^) in CRPS, with expansions in effector Th (CD62L^−^CD45RA^+^) and effector memory Th subsets (CD62L^−^CD45RA^−^). **(B)** Naive Tregs (CD4^+^CD25^+^CD127^−^ FOXP3^+^CD62L^+^CD45RA^+^) were also decreased in CRPS. Boxplots show IQR, median (solid lines), and mean (+) for each T cell subset, with whiskers extending to 1.5x IQR. Each point represents an individual participant. Statistical comparisons were performed using Welch’s t-tests for parametric data (determined by Shapiro-Wilk test) or Mann-Whitney U-tests for non-parametric data, with *P < 0.05; **P < 0.01. Statistical outliers (grey points) were excluded using ROUT (Q = 1%). **(C)** Two helper T cell metaclusters (naive and Temra) showed higher expression of pro-inflammatory signalling molecules. Heatmap shows log-fold changes in activity marker expression for CD3^+^ FlowSOM metaclusters between CRPS and controls, with a ratio of 0 (white) indicating equal expression, a ratio greater than 0 (orange/red) indicating CRPS>control, and a ratio less than 0 (blue) indicating CRPS<control, with *P_adj_ < 0.05.

Differential expression analysis performed on CD3^+^ metaclusters revealed some changes in T cell inflammation and activation marker expression between CRPS and control groups (Fig. 4C). Naive Th cells upregulated p65 NFκB in CRPS (P_adj_ < 0.0001), and a Temra helper subset (CD4^+^CD62L^lo^CD45RA^+^CD45RO^+^) showed increased CXCR3 expression (P_adj_ = 0.0334). Altogether, T cell subset activity in CRPS appeared largely similar to controls, apart from differences in inflammatory marker expression displayed by some helper T cell subsets.

### CRPS participants had altered serum expression of immunoglobulin variants and increased levels of proteins associated with systemic inflammation

SWATH MS-based proteomics identified 14 serum proteins that were significantly differentially expressed in CRPS. (Figure 5A; Table 2), most of which are implicated in the innate or adaptive immune system. Several inflammatory mediators were upregulated in the CRPS group, including C-reactive protein (CRP), cysteine-rich secretory protein 3 (CRIS3), and ficolin-3 (FCN3). Downregulated proteins were mostly immunoglobulin variants, including two heavy-chain (HV338 and HV5X1) and four lambda variants (LV861, LV151, LV473, LV949). GSEA of the differentially expressed proteins revealed involvement in immune regulation and activation processes, including the lectin pathway of complement activation and multiple pattern recognition receptor signalling pathways (Fig. 5B), and an association with autoimmune disease terms from the Jensen DISEASE library (Fig. 5C). Reactome pathway analysis confirmed involvement in complement pathways, including initial triggering of complement, classical antibody-mediated complement activation, and creation of C2 and C4 activators (Fig. 5D-E). Several pathways associated with BCR activation and Fc-receptor signalling were enriched among the differentially expressed heavy-chain and lambda Ig variants (Fig. 5D), although their downregulation more likely reflects an altered serum antibody repertoire in CRPS than a change in these specific signalling pathways. Further changes seen in the CRPS group were differential expression of coagulation factors (FA5, FA11) and apolipoproteins (APOC4, APOF), possibly signalling disruption to the clotting cascade and pathways of lipoprotein assembly, remodelling, and clearance (Fig. 5D-E). Overall, proteomic analysis identified changes in serum antibody expression in CRPS, and increased levels of circulating proteins associated with systemic inflammation.

**Figure 5.**
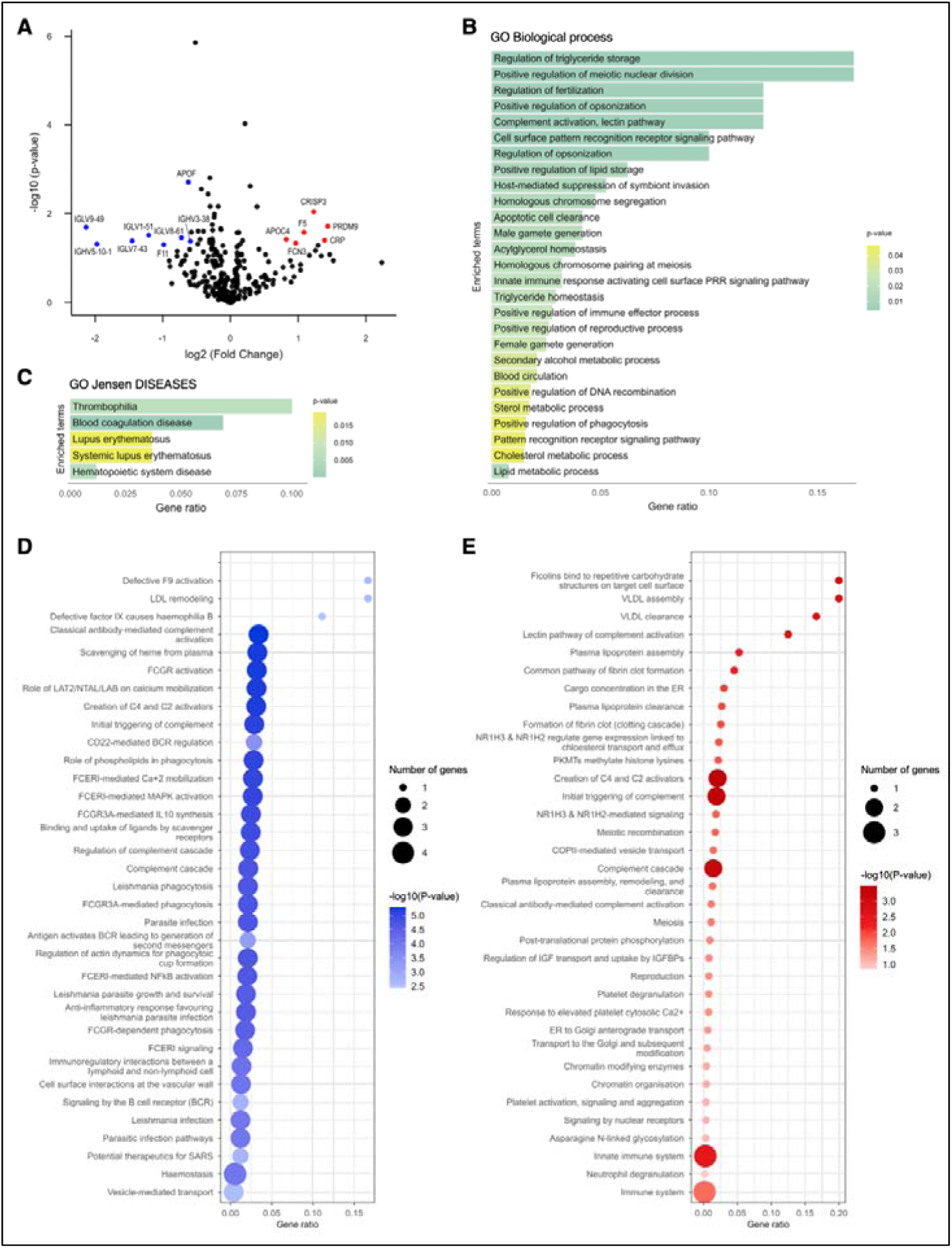
Differences in serum protein expression between CRPS subjects (*n* = 42) and healthy donors (*n* = 20) revealed by SWATH-MS proteomic analysis. **(A)** Volcano plot shows proteins that were differentially expressed in CRPS compared to controls. Positive log2(FC) values indicate upregulation in CRPS, and negative values represent downregulation, with significantly increased or decreased proteins (FC > ±1.5, P < 0.05) shown in red or blue respectively. (**B-C**) Bar plots show GO biological process and DISEASE database terms that were significantly enriched (P_adj_ < 0.05) among the altered proteins, with bar colour representing P_adj_ for the enriched term. (**D-E**) Dot plots show the top 30 Reactome pathways (all P_adj_ < 0.05) that were enriched among the significantly downregulated (panel **D**) and upregulated proteins (panel **E**) in CRPS. Point gradient represents -log10(P_adj_), and point size represents the number of proteins enriched in the relevant pathway.

**Table 2.** Differentially expressed serum proteins between CRPS subjects and healthy donors identified by SWATH MS-based proteomics.

| **Protein** | **Fold change** | **P-value** | **Gene name** | **Protein description** |
|---|---|---|---|---|
| PRDM9 ↑ | 2.7079 | 0.0192* | PRDM9 | Histone-lysine N-methyltransferase PRDM9 |
| CRP ↑ | 2.6212 | 0.0395* | CRP | C-reactive protein |
| CRIS3 ↑ | 2.3448 | 0.0090** | CRISP3 | Cysteine-rich secretory protein 3 |
| FA5 ↑ | 2.1223 | 0.0261* | F5 | Coagulation factor V |
| FCN3 ↑ | 1.9502 | 0.0459* | FCN3 | Ficolin-3 |
| APOC4 ↑ | 1.7696 | 0.0378* | APOC4 | Apolipoprotein C-IV |
| HV338 ↓ | 0.6642 | 0.0417* | IGHV3-38 | Immunoglobulin heavy variable 3-38 |
| APOF ↓ | 0.6498 | 0.0020** | APOF | Apolipoprotein F |
| LV861 ↓ | 0.6058 | 0.0348* | IGLV8-61 | Immunoglobulin lambda variable 8-61 |
| FA11 ↓ | 0.5052 | 0.0499* | F11 | Coagulation factor XI |
| LV151 ↓ | 0.4335 | 0.0303* | IGHV1-51 | Immunoglobulin lambda variable 1-51 |
| LV743 ↓ | 0.3657 | 0.0408* | IGLV7-43 | Immunoglobulin lambda variable 7-43 |
| HV5X1 ↓ | 0.2546 | 0.0481* | IGHV5-10-1 | Immunoglobulin heavy variable 5-10-1 |
| LV949 ↓ | 0.2280 | 0.0202* | IGLV9-49 | Immunoglobulin lambda variable 9-49 |

Table shows fold change, P-value, gene name and description for proteins that were differentially expressed in the serum of CRPS subjects (*n* = 42) compared to healthy controls (*n* = 20), with a fold change > ±1.5 and P < 0.05 (unpaired t-test). Arrows indicate whether a given protein was significantly upregulated or downregulated in CRPS. \*P < 0.05; \*\*P < 0.01.

### CRPS participants had altered IgA and IgG autoantibody profiles against a small number of native antigen targets

Penetrance fold change (pFC) analysis of serum IgG and IgA autoantibodies against human antigens from the iOme^®^ Discovery Protein Array revealed some differences in autoantibody profiles between CRPS and healthy control groups (Figure 6). CRPS participants had increased IgA autoantibodies against nine proteins (Table 3; Fig. 6A; Supplementary Figure 11B), including SH3GL1 (pFC = 3; P = 0.0055) and STK25 (pFC = 5, P = 0.045). The CRPS cohort also had increased IgG autoantibodies against the protein LRRFIP2 (pFC = 5; P = 0.071), with modest increases seen for three other proteins (Table 3; Fig. 6B; Supplementary Fig. 11A). Surprisingly, the levels of all other IgG autoantibodies identified by pFC analysis were lower in CRPS compared to the control group. Overall, there was a high degree of individual variation in CRPS autoantibody profiles, and no antigen targets were consistently increased across the whole patient cohort.

**Figure 6.**
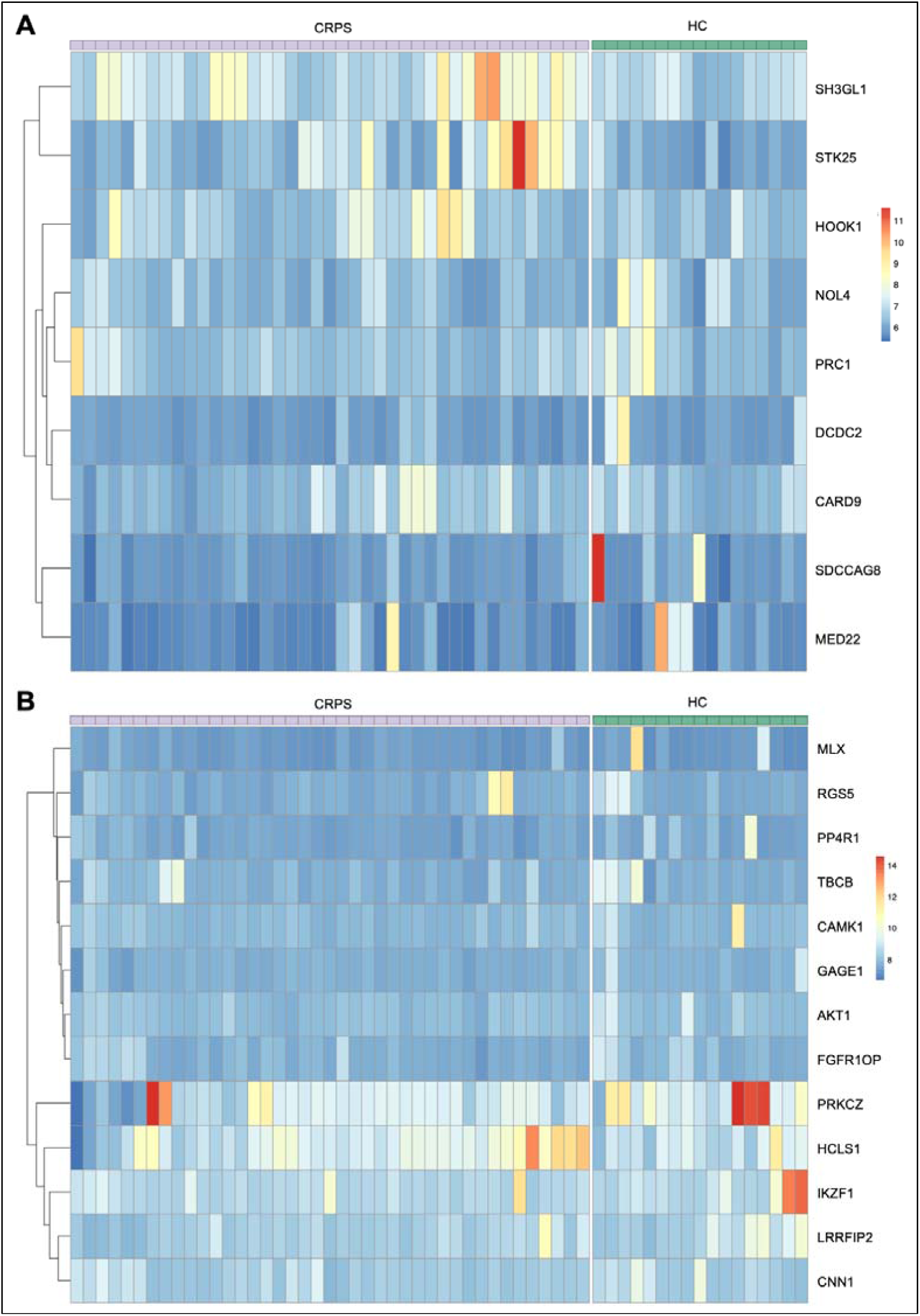
Serum autoantibody profiles against top antigens identified by penetrance fold change (pFC) analysis of CRPS subjects (*n* = 42) and healthy donors (*n* = 18) using the iOme^®^ Discovery Protein Array. Heatmaps represent log2-transformed normalised values for **(A)** IgA and **(B)** IgG autoantibodies across all samples against the top antigens identified using pFC analysis of CRPS vs. controls, with pFC ≥ 2 and P < 0.2 (Fisher’s exact test).

**Table 3.** Top antigens from the iOme^®^ Discovery Protein Array identified by penetrance fold change analysis of serum autoantibody profiles between CRPS subjects and healthy donors.

| Antigen | pFC (CRPS) | pFC (HC) | P-value | Antigen name |
|---|---|---|---|---|
| *IgA autoantibodies* |  |  |  |  |
| SH3GL1 | 3 | 0 | 0.0055* | SH3 domain containing GRB2 like 1 |
| DCDC2 | 0 | 3 | 0.022* | Doublecortin domain containing 2 |
| STK25 | 5 | 2 | 0.045* | Serine/threonine kinase 25 |
| NOL4 | 0 | 4 | 0.082* | Nucleolar protein 4 |
| SDCCAG8 | 0 | 28 | 0.082* | SHH signalling/ciliogenesis regulator SDCCAG8 |
| HOOK1 | 3 | 0 | 0.093* | Hook microtubule tethering protein 1 |
| MED22 | 5 | 3 | 0.14 | Mediator complex subunit 22 |
| PRC1 | 5 | 3 | 0.14 | Protein regulator of cytokinesis 1 |
| CARD9 | 3 | 0 | 0.17 | Caspase recruitment domain family member 9 |
| *IgG autoantibodies* |  |  |  |  |
| LRRFIP2 | 5 | 3 | 0.071* | LRR-binding FLII interacting protein 2 |
| AKT1 | 0 | 2 | 0.082* | AKT serine/threonine kinase 1 |
| CAMK1 | 0 | 6 | 0.082* | Calcium/calmodulin-dependent protein kinase 1 |
| CNN1 | 0 | 3 | 0.082* | Calponin 1 |
| FGFR1OP | 0 | 2 | 0.082* | Centrosomal protein 43 |
| GAGE1 | 0 | 2 | 0.082* | G antigen 1 (cancer/testis antigen 4.1) |
| MLX | 0 | 12 | 0.082* | MAX dimerization protein MLX |
| PPP4R1 | 0 | 4 | 0.082* | Protein phosphatase 4 regulatory subunit 1 |
| PRKCZ | 8 | 25 | 0.11* | Protein kinase C zeta |
| IKZF1 | 6 | 25 | 0.14* | IKAROS family zinc finger protein 1 |
| RGS5 | 11 | 3 | 0.14* | Regulator of G-protein signalling 5 |
| TBCB | 4 | 3 | 0.14* | Tubulin folding fofactor B |
| HCLS1 | 3 | 5 | 0.15* | Hematopoietic cell-specific lyn substrate 1 |

Table shows penetrance fold change (pFC), P-value, and full name of the top antigens identified by pFC analysis of serum IgA and IgG autoantibodies between CRPS and healthy control groups, with FC ≥ 2 and P < 0.2 (Fisher’s exact test).

## Discussion

This study used a high-dimensional immunophenotyping approach to explore changes in circulating B cell and autoantibody profiles in people with complex regional pain syndrome. Mass cytometry findings identified enrichment in autoimmune-associated heavy-chain gene segments among class-switched B cells in CRPS participants, who also showed shifts in the memory B cell repertoire away from resting-state subsets and towards atypical/double-negative memory phenotypes. Together, these results are suggestive of an extrafollicular B cell response in CRPS that could enable production of pathogenic, innate-like autoantibodies, contributing to sustained autoinflammation and pain. Further changes in serum protein expression and autoantibody profiles are reflective of persistent immune dysfunction in CRPS that could also contribute to autoinflammation. Collectively, these findings support a role for dysregulated B cell tolerance in the onset and maintenance of pathogenic autoimmune responses in CRPS.

### CRPS participant memory B cell profiles resemble those seen in classical autoimmune disease to suggest increased extrafollicular B cell activation

CSM B cells in a resting/inactive state were reduced in CRPS, without changes in the proportions of transitional, mature naive, USM, MZ, or antibody-secreting B cell subsets. Similar shifts away from the resting memory pool are observed in prototypical autoimmune conditions such as SLE and RA, where patients show expansions in activated switched memory B cell populations in line with disease activity and post-treatment relapse [59–63]. Memory B cells have a central role in autoimmune diseases such as these, where their naturally low reactivation thresholds, high proliferative capacity, and prolonged survival can facilitate persistent self-directed immune activation and autoantibody production [64–66]. The reduction in resting memory B cells seen in CRPS therefore indicates a shift in the memory B cell compartment that could reflect a state of ongoing immune activation. A central characteristic of memory B cells is their rapid proliferation and differentiation into ASCs upon re-exposure to a previously encountered antigen [64, 65]. This can proceed by helper T cell-mediated processes in the germinal centre (GC), where somatic hypermutation (SHM) and affinity selection lead to the generation of long-lasting ASCs with high antigen affinity, but can also occur via T cell-independent extrafollicular pathways, by which reactivated memory cells differentiate into short-lived, lower-affinity plasmablasts without re-entering the GC [65]. The extrafollicular region has been identified as key site for tolerance breakdown and autoantibody production in autoimmune disease, with non-GC responses driving the expansion of poorly mutated, autoreactive ASCs in mice that can lead to systemic autoimmunity [67–69]. In this study, evidence of an extrafollicular response is offered by the increased proportions of distinct ‘double-negative’ (CD27^−^IgD^−^) memory subsets in CRPS: DN2 B cells (CD38^−^CD21^−^), FCRL4^+^ DN2 B cells, and a DN2-like subset with low levels of CD38. DN2 B cells are also expanded in SLE and RA, displaying enriched autoreactivity compared to classical CSM B cells and correlating with serum autoantibody levels [55, 70–72], which has led to the suggestion that they function as extrafollicular-derived ASC precursors [55, 73–75]. Together with the decrease in resting CSM B cells, this indicates increased extrafollicular activation in people with persistent CRPS, shifting the B cell pool away from classical memory phenotypes and towards atypical, low-affinity, potentially pathogenic DN B cell subpopulations.

DN B cells are also expanded in contexts of chronic antigen stimulation and immune exhaustion, such as HIV, malaria, and cancer (along with autoimmune disease) [69, 57, 76, 77]. CRPS subjects had a higher proportion of DN2 B cells expressing FCRL4 (Fc-receptor like protein 4), an inhibitory molecule that can signal exhaustion when upregulated on circulating B cells [78–80]. In RA, synovial FCRL4^+^ B cells also overexpress autoreactive receptors and produce anti-citrullinated autoantibodies [80, 81], supporting a pathogenic role for the expanded FCRL4^+^ DN2 population observed in CRPS. Unsupervised B cell clustering also revealed a CD21^lo^CD20^lo^ DN subset which upregulated CD95, PD-1 and PD-L1 in CRPS, consistent with an exhausted B cell phenotype [82, 83]. CD95 and PD-1/PD-L1 signalling also regulate GC tolerance, and expansions in CD95^+^ and PD-L1^+^ B cells correlate with follicular helper T cell (Tfh) abundance and disease activity in SLE [84–86, 63], so these findings could indicate disturbances to the regulation of T cell-mediated B cell maturation as well as extrafollicular pathways. Further impairment to B cell regulation in CRPS could derive from altered expression of CD22, an inhibitory co-receptor that helps maintain self-tolerance by regulating BCR activation thresholds [87, 88]. CD22 was downregulated by two CD19^+^ populations in CRPS, and across the broader B cell compartment in the Melbourne CRPS cohort. Defects in CD22 signalling are associated with RA, SLE, and type 1 diabetes, and can promote autoantibody production and systemic autoimmunity in mice [89–94], so the current evidence warrants further investigation into altered CD22 activity as a mechanism of dysfunctional self-tolerance and autoimmunity in CRPS. Overall, these changes in classical and atypical memory B cell populations in CRPS are suggestive of dysregulated B cell maturation, which may lead to the proliferation of autoreactive clones.

Further evidence of altered B cell maturation in CRPS derives from the observed increases in the expression of autoimmune-associated gene segment by CSM B cells, as indicated by higher G6 and 9G4 binding in this compartment. The G6 and 9G4 antibodies bind cross-reactive idiotypes (CRI) found on Ig heavy-chains encoded by the VH1-69 and VH4-34 gene segments, respectively, which are overexpressed by B cells in autoimmune disease, contributing to autoantibody secretion [95–97]. IGHV1-69 is one of several germline genes encoding rheumatoid factor autoantibodies, which bind the Fc region of IgG and contribute to immunocomplex formation in RA, driving the distinctive joint inflammation and synovial tissue damage [97, 98]. IGHV4-34 autoantibodies primarily recognise carbohydrate self-epitopes displayed by erythrocytes and B lymphocytes, and represent a proportion of the disease-characteristic anti-dsDNA antibodies seen in SLE [99–106]. IGHV1-69 and IGHV4-34 B cells are also present in healthy people, but are not usually detected among class-switched B cells due to somatic hypermutation, which introduces mutations in the G6 and 9G4 idiotypes that reduce autoreactivity and increase affinity for foreign antigen, thus preventing G6 and 9G4 binding [97, 107]. In this study, only the CRPS group showed high levels of G6 and 9G4 binding among CSM B cell subsets, evidence of unmutated IGHV1-69 and IGHV4-34 in this compartment. This is further indication of an extrafollicular memory response in CRPS, as similar observations of poorly mutated B cell clones and autoantibodies in people with RA, Sjögren’s syndrome, or SLE have led to the suggestion that a significant proportion of circulating autoantibodies in these conditions have an extrafollicular origin [68, 108, 109].

The increased 9G4 binding seen here in the IgD^+^CD27^+^ unswitched memory (USM) compartment of patients is particularly strong evidence of an extrafollicular response in CRPS. USM B cells function as an innate-like memory subset, differentiating via extrafollicular pathways into short-lived plasmablasts that secrete low-affinity IgM [65, 110, 111]. Accordingly, these cells show enriched autoreactivity in SLE, where the frequency of circulating IGHV4-34^+^IgM^+^ USM B cells correlates with patient symptoms, disease activity, and serum complement levels [112]. IgM autoantibodies can activate both the classical and alternative complement pathways, so taken with the current findings, this could implicate extrafollicular-derived, innate-like memory B cell populations as a source of the autoreactive IgM that contributes to pain via complement activation in the fracture/cast rodent model of CRPS [113, 114, 12]. Further, our findings indicate this mechanism could be particularly significant in women with CRPS, as female patients showed higher 9G4 binding in the USM and CSM compartments compared to male patients. This is consistent with prior observations that healthy women have higher baseline IgM levels than men [116], but also that autoantibody-mediated autoimmune conditions are generally more prevalent in women, suggestive of sex differences in the pathophysiology of autoimmune responses (as reviewed in [117]). Rodent models have also identified evidence of sexual dimorphism in autoimmune-mediated mechanisms of post-fracture CRPS pain and neuropathic pain specifically [118–120]. Together, these observations warrant further enquiry into sex-specific mechanisms of autoantibody-mediated pain in CRPS.

This breakdown in peripheral tolerance in CRPS could be induced by the early-stage inflammatory environment caused by the inciting tissue trauma [109]. This would align with observations that CRPS patient autoantibodies only recapitulate disease symptoms in rodents that have a hindlimb injury (unlike in fibromyalgia, where autoantibody transfer alone is sufficient to induce pain behaviours in recipient mice) [121, 122], which established the ‘two-hit’ model of CRPS pain: pre-existing innate autoantibodies (the ‘first hit’) become pathogenic after the inciting tissue trauma (the ‘second hit’) [2, 12]. The current findings indicate that this occurs when extrafollicular responses are triggered by trauma-induced inflammatory activation, leading to the production of low-affinity pathogenic autoantibodies that contribute to long-term autoinflammation and pain.

### CRPS is associated with expansions in antigen-experienced helper T cell populations and enhanced inflammatory T cell signalling

CRPS participants in this study had a reduced naive helper T cell compartment and expanded effector and effector memory Th populations. This general shift from naive to antigen-experienced T cell phenotypes is consistent with our prior findings [24], although we previously saw increases in CD4^+^ and CD8a^+^ central memory T cells in CRPS. This may reflect the use of alternative markers to define central memory T cells (CD62L versus CCR7), and the comparison of proportional abundance rather than absolute cell counts. In the present study, naive Th cells also showed higher phosphorylated p65 expression, indicating enhanced NFκB signalling, which controls pro-inflammatory T cell activation and cytokine production, induces Th1 differentiation, and is also upregulated by circulating T cell populations in both CRPS and RA [24, 123–126]. Another driver of Th1 differentiation, CXCR3 [126], was also increased in CRPS, in the Temra helper cell population. Together with previous reports of elevated tumour necrosis factor (TNF), IL-2, and soluble IL-2 receptor (sIL-2R) expression, and increased T-bet^+^ T cell frequency in CRPS [128, 27, 24], these findings support a Th1 contribution to chronic inflammation during the condition.

Unlike our earlier study [24], no further increases in inflammatory T cell markers were detected, which may reflect differences in cohort composition. The current study included nearly twice as many participants as the earlier study, which also had a higher proportion of acute CRPS subjects compared to the present cohort, where all but two patients were more than 6 months post-injury. This aligns with existing evidence that early-stage inflammatory activation in CRPS diminishes over time and becomes superseded by pathogenic adaptive immune responses [22, 129, 11]. This could be facilitated by the anti-inflammatory, immunosuppressive, anti-nociceptive activity of Treg cells [130–132], which showed a reduced naive population in this study, consistent with previous findings of Treg expansion in CRPS [24, 25] (although these cells may be insufficiently activated or non-functional in CRPS to enable persistent dysregulation of other immune cell populations). Further research into the role of Tregs and their therapeutic potential in CRPS is required. Overall, these results implicate helper T cell dysfunction in chronic CRPS, through both effector and effector memory responses and impaired immune regulation.

### Chronic CRPS patients show increased serum expression of complement activators

CRPS patients showed increased levels of circulating C-reactive protein (CRP) and ficolin-3 (FCN3), both of which are pattern recognition molecules involved in complement activation. CRP is an acute-phase protein that triggers the classical complement pathway by interacting with Fc-receptors [133, 134], while FCN3 is a potent stimulator of the lectin pathway of complement activation [135]. Typically, increases in CRP expression (>3-fold) are only observed directly after acute infection or injury [136], so the maintained 2-fold upregulation of this protein in the current chronic CRPS cohort is strong evidence of ongoing low-level inflammation that is pathogenic and not merely associated with tissue trauma. CRP and FCN3 are also implicated in autoantibody-mediated autoimmune conditions. CRP, which can recognise self-antigens via pattern recognition, is increased in serum during RA and correlates with disease activity and tissue damage [137], while increased FCN3 expression in SLE is associated with serum autoantibody enrichment [138]. Importantly, complement activation via complement receptors on dermal macrophages and spinal microglia is thought to represent a central mechanism of CRPS-like pain in the fracture/cast rodent model [114, 139]. Moreover, complement receptor expression on osteoblasts has been shown to impair fracture healing [140], which may also attenuate normal healing in CRPS. Further changes that could signal long-term immune dysregulation were the reductions in multiple IgH and IgL proteins in CRPS, which could reflect a shift in the circulating antibody repertoire towards autoreactive variants like IGHV1-69 and IGHV4-34, although this should be confirmed through further characterisation of the BCR repertoire in CRPS. Together with mass cytometry findings, the results of serum proteomic analysis offer evidence for pathogenic antibody-mediated complement activation in clinical CRPS.

### The diversity of CRPS autoantibody profiles may reflect its heterogenous presentation

Autoantibody profiling revealed a small number of antigen targets that were enriched among serum IgG and IgA autoantibody profiles in CRPS. People with CRPS had significantly higher IgA autoantibody levels against SH3GL1 and STK25, both of which have been implicated in autoimmune disease pathogenesis. SH3GL1 encodes an endocytic protein called endophilin A2, and may have a regulatory, protective role in the context of autoimmune arthritis, as defective expression of this gene in rodents is observed to alter effector T cell activity and activation thresholds for autoreactive T cells [141]. It is also upregulated by CD4^+^ T cell populations in RA patients, where it can modulate TCR signalling [141]. Conversely, STK25 (serine/threonine protein kinase 25) has been identified as a positive regulator of inflammatory cytokine production and is overactive in people with SLE [142]. These proteins may therefore be of interest for future investigation into the targets of autoreactive CRPS antibodies involved in inflammation and autoimmunity, although they should first be validated by an alternate method. The role of IgA, which normally supports mucosal and barrier immunity, is poorly understood in the context of CRPS, as existing evidence is primarily concerned with pathogenic IgG and IgM. RA autoantibodies (including rheumatoid factors) can be IgA-switched, which may predict more severe disease [143, 144], so the current findings warrant further investigation into IgA autoantibodies in CRPS. Together with previous autoantibody studies [14–21], these results also indicate a high degree of patient stratification in the autoantibody targets of CRPS, possibly linked to disease presentation or clinical features such as triggering injury. Higher-powered studies are required to understand the full landscape of autoreactive antibodies involved in CRPS.

### Conclusions

This study identified several changes in the circulating B cell repertoires of people with CRPS consistent with chronic autoimmune activity. The increased autoreactive idiotype usage in the class-switched B cell compartment offers evidence of an extrafollicular response that leads to impaired peripheral tolerance in CRPS. Along with the changes in memory B cell subsets, helper T cell activity, and autoantibody profiles, these findings support the view that persistent CRPS is mediated by pathogenic adaptive immune activity, including self-directed antibody responses. Additional autoinflammatory contributions are highlighted by the increased expression of complement activators in CRPS. Further investigation is required to understand how tolerance breakdown occurs in this condition, and to confirm the identity of pain-promoting autoantibodies. This will provide a deeper understanding of how pain is established and maintained in CRPS, and therefore how it can be addressed in a targeted and effective way.

### Limitations

This study has some limitations that should be addressed by future studies. Our findings demonstrate an association between B cell dysregulation and CRPS, not causation, so further investigation into their relationship with broader mechanisms of CRPS is required. Proteomic and autoantibody findings should be validated in more diverse cohorts with a larger control sample, allowing for better group comparison and stratification by patient demographics and disease features. This would also help determine the clinical relevance of the serum proteins and autoantibody targets identified. Including more male participants in these investigations would allow for a better understanding of sex differences in autoimmune mechanisms of CRPS. Finally, a longitudinal study would be beneficial in understanding how these immune profiles evolve from the acute to chronic stage. Together, these studies would help to clarify the precise relationship between peripheral autoimmune activity and regional pain in CRPS.

## Author contributions

**ACM** designed and optimised both mass cytometry experiments, stained and acquired blood samples, performed manual gating and unsupervised analysis of mass cytometry data, conducted downstream analysis of proteomic data, produced the figures, wrote the first draft of the manuscript, and edited the manuscript; **FAC** assisted with mass cytometry experiments and data analysis; **JFK** assisted with mass cytometry data analysis; **BH, SBA** and **SK** acquired and processed proteomic data, and **SBA** wrote the corresponding supplementary methods; **JAO** contributed to mass cytometry panel optimisation and FlowSOM analysis; **YH** recruited participants and processed clinical data; **RW** collected clinical data, took blood samples and performed initial blood processing; **VM** collected clinical data; **MAR**, **NC**, and **PG** assisted with participant recruitment; **JHR** contributed to with mass cytometry panel design, supervised data interpretation, and edited the manuscript; **PJA** conceived of, supervised and procured funding for the study, assisted with mass cytometry experiments, and edited the manuscript. All authors read and approved the final manuscript.

## Acknowledgements

We are first grateful to all the volunteers and their families whose participation made this study possible. Further thanks to the Sydney Cytometry Core Research Facility (University of Sydney and Centenary Institute, Sydney, NSW, Australia) for providing equipment, reagents, and technical assistance with mass cytometry experiments, and to Houston Omics Collaborative at the University of Houston (Houston, USA) who ran the iOme^®^ Protein Array and performed corresponding data analysis.

## References

1 Lee, D. H., Noh, E. C., Kim, Y. C., Hwang, J. Y., Kim, S. N., Jang, J. H., Byun, M. S & Kang, D. H. 2014. Risk Factors for Suicidal Ideation among Patients with Complex Regional Pain Syndrome. Psychiatry Investig, 11, 32–8.

2 Goebel, A., Andersson, D., Helyes, Z., Clark, J. D., Dulake, D. & Svensson, C. 2022. The autoimmune aetiology of unexplained chronic pain. Autoimmun Rev, 21, 103015.

3 Bruehl, S. 2015. Complex regional pain syndrome. BMJ, 351, h2730.

4 Sharma, A., Agarwal, S., Broatch, J. & Raja, S. N. 2009. A web-based cross-sectional epidemiological survey of complex regional pain syndrome. Reg Anesth Pain Med, 34, 110–5.

5 Lohnberg, J. A. & Altmaier, E. M. 2013. A Review of Psychosocial Factors in Complex Regional Pain Syndrome. J Clin Psychol Med Settings, 20, 247–254.

6 Ferraro, M. C., Cashin, A. G., Wand, B. M., Smart, K. M., Berryman, C., Marston, L., Moseley, G. L., Mcauley, J. H. & O’connell, N. E. 2023. Interventions for treating pain and disability in adults with complex regional pain syndrome-an overview of systematic reviews. Cochrane Database Syst Rev, 6, CD009416.

7 Goebel, A. & Blaes, F. 2013. Complex regional pain syndrome, prototype of a novel kind of autoimmune disease. Autoimmun Rev, 12, 682–6.

8 Tekus, V., Hajna, Z., Borbely, E., Markovics, A., Bagoly, T., Szolcsanyi, J., Thompson, V., Kemeny, A., Helyes, Z. & Goebel, A. 2014. A CRPS-IgG-transfer-trauma model reproducing inflammatory and positive sensory signs associated with complex regional pain syndrome. Pain, 155, 299–308.

9 Cuhadar, U., Gentry, C., Vastani, N., Sensi, S., Bevan, S., Goebel, A. & Andersson, D. A. 2019. Autoantibodies produce pain in complex regional pain syndrome by sensitizing nociceptors. Pain, 160, 2855–2865.

10 Helyes, Z., Tekus, V., Szentes, N., Pohoczky, K., Botz, B., Kiss, T., Kemeny, A., Kornyei, Z., Toth, K., Lenart, N., Abraham, H., Pinteaux, E., Francis, S., Sensi, S., Denes, A. & Goebel, A. 2019. Transfer of complex regional pain syndrome to mice via human autoantibodies is mediated by interleukin-1-induced mechanisms. Proc Natl Acad Sci U S A, 116, 13067–13076.

11 Guo, T. Z., Shi, X., Li, W. W., Wei, T., Clark, J. D. & Kingery, W. S. 2017. Passive transfer autoimmunity in a mouse model of complex regional pain syndrome. Pain, 158, 2410–2421.

12 Guo, T. Z., Wei, T., Tajerian, M., Clark, J. D., Birklein, F., Goebel, A., Li, W. W., Sahbaie, P., Escolano, F. L., Herrnberger, M., Kramer, H. H. & Kingery, W. S. 2020. Complex regional pain syndrome patient immunoglobulin M has pronociceptive effects in the skin and spinal cord of tibia fracture mice. Pain, 161, 797–809.

13 Rose, N. R. & Bona, C. 1993. Defining criteria for autoimmune diseases (Witebsky’s postulates revisited). Immunol Today, 14, 426–30.

14 Blaes, F., Schmitz, K., Tschernatsch, M., Kaps, M., Krasenbrink, I., Hempelmann, G. & Brau, M. E. 2004. Autoimmune etiology of complex regional pain syndrome (M. Sudeck). Neurology, 63, 1734–6.

15 Kohr, D., Tschernatsch, M., Schmitz, K., Singh, P., Kaps, M., Schafer, K. H., Diener, M., Mathies, J., Matz, O., Kummer, W., Maihofner, C., Fritz, T., Birklein, F. & Blaes, F. 2009. Autoantibodies in complex regional pain syndrome bind to a differentiation-dependent neuronal surface autoantigen. Pain, 143, 246–251.

16 Kohr, D., Singh, P., Tschernatsch, M., Kaps, M., Pouokam, E., Diener, M., Kummer, W., Birklein, F., Vincent, A., Goebel, A., Wallukat, G. & Blaes, F. 2011. Autoimmunity against the beta2 adrenergic receptor and muscarinic-2 receptor in complex regional pain syndrome. Pain, 152, 2690–2700.

17 Dubuis, E., Thompson, V., Leite, M. I., Blaes, F., Maihofner, C., Greensmith, D., Vincent, A., Shenker, N., Kuttikat, A., Leuwer, M. & Goebel, A. 2014. Longstanding complex regional pain syndrome is associated with activating autoantibodies against alpha-1a adrenoceptors. Pain, 155, 2408–17.

18 Dirckx, M., Schreurs, M. W., DE Mos, M., Stronks, D. L. & Huygen, F. J. 2015. The prevalence of autoantibodies in complex regional pain syndrome type I. Mediators Inflamm, 2015, 718201.

19 Shubayev, V. I., Strongin, A. Y. & Yaksh, T. L. 2018. Structural homology of myelin basic protein and muscarinic acetylcholine receptor: Significance in the pathogenesis of complex regional pain syndrome. Mol Pain, 14, 1744806918815005.

20 Baerlecken, N. T., Gaulke, R., Pursche, N., Witte, T., Karst, M. & Bernateck, M. 2019. Autoantibodies against P29ING4 are associated with complex regional pain syndrome. Immunol Res, 67, 461–468.

21 VAN DER Spek, D. P. C., Hoffenkamp, R. H., Huygen, F., Bharwani, K. D., Eijkelkamp, N. & Dirckx, M. 2026. Autonomic Receptor Autoantibodies in Complex Regional Pain Syndrome and Other Chronic Pain Conditions: A Cross-Sectional Analysis. Biomedicines, 14.

22 Parkitny, L., Mcauley, J. H., DI Pietro, F., Stanton, T. R., O’connell, N. E., Marinus, J., VAN Hilten, J. J. & Moseley, G. L. 2013. Inflammation in complex regional pain syndrome: a systematic review and meta-analysis. Neurology, 80, 106–17.

23 Schlereth, T., Drummond, P. D. & Birklein, F. 2014. Inflammation in CRPS: role of the sympathetic supply. Auton Neurosci, 182, 102–7.

24 Russo, M. A., Fiore, N. T., VAN Vreden, C., Bailey, D., Santarelli, D. M., Mcguire, H. M., FAZEKAS DE ST Groth, B. & Austin, P. J. 2019. Expansion and activation of distinct central memory T lymphocyte subsets in complex regional pain syndrome. J Neuroinflammation, 16, 63.

25 Heyn, J., Azad, S. C. & Luchting, B. 2019. Altered regulation of the T-cell system in patients with CRPS. Inflamm Res, 68, 1–6.

26 Kaufmann, I., Eisner, C., Richter, P., Huge, V., Beyer, A., Chouker, A., Schelling, G. & Thiel, M. 2007. Lymphocyte subsets and the role of TH1/TH2 balance in stressed chronic pain patients. Neuroimmunomodulation, 14, 272–80.

27 Bharwani, K. D., Dirckx, M., Stronks, D. L., Dik, W. A., Schreurs, M. W. J. & Huygen, F. 2017. Elevated Plasma Levels of sIL-2R in Complex Regional Pain Syndrome: A Pathogenic Role for T-Lymphocytes? Mediators Inflamm, 2017, 2764261.

28 Li, W. W., Yang, Y., Guo, T. Z., Sahbaie, P., Shi, X. Y., Guang, Q., Kingery, W. S., Herzenberg, L. A. & Clark, J. D. 2021. IL-6 signaling mediates the germinal center response, IgM production and nociceptive sensitization in male mice after tibia fracture. Brain Behav Immun, 94, 148–158.

29 Lacagnina, M. J., Willcox, K. F., Boukelmoune, N., Bavencoffe, A., Sankaranarayanan, I., Barratt, D. T., Zuberi, Y. A., Dayani, D., Chavez, M. V., Lu, J. T., Farinotti, A. B., Shiers, S., Barry, A. M., Mwirigi, J. M., Tavares-Ferreira, D., Funk, G. A., Cervantes, A. M., Svensson, C. I., Walters, E. T., Hutchinson, M. R., Heijnen, C. J., Price, T. J., Fiore, N. T. & Grace, P. M. 2024. B cells drive neuropathic pain-related behaviors in mice through IgG-Fc gamma receptor signaling. Sci Transl Med, 16, eadj1277.

30 Fiore, N. T., Willcox, K. F., Grieco, A. R., Dayani, D., Zuberi, Y. A., Heijnen, C. J. & Grace, P. M. 2025. Autoreactive immunoglobulin G levels and Fc receptor gamma subunit upregulation drive mechanical allodynia after nerve constriction or crush injury. Pain, 166, 2804–2817.

31 Harden, R. N., Bruehl, S., Stanton-Hicks, M. & Wilson, P. R. 2007. Proposed new diagnostic criteria for complex regional pain syndrome. Pain Med, 8, 326–31.

32 Dworkin, R. H., Turk, D. C., Revicki, D. A., Harding, G., Coyne, K. S., Peirce-Sandner, S., Bhagwat, D., Everton, D., Burke, L. B., Cowan, P., Farrar, J. T., Hertz, S., Max, M. B., Rappaport, B. A. & Melzack, R. 2009. Development and initial validation of an expanded and revised version of the Short-form McGill Pain Questionnaire (SF-MPQ-2). Pain, 144, 35–42.

33 Lovibond, P. F. & Lovibond, S. H. 1995. The structure of negative emotional states: comparison of the Depression Anxiety Stress Scales (DASS) with the Beck Depression and Anxiety Inventories. Behav Res Ther, 33, 335–43.

34 Sullivan, M. J. L., Bishop, S. R. & Pivik, J. 1995. The Pain Catastrophizing Scale: Development and validation. Psychol Assess, 7, 524–532.

35 Miller, R., Kori, S., & Todd, D. 1991. The Tampa Scale: a Measure of Kinesiophobia. Clin J Pain, 7, 51.

36 Nicholas, M. K. 2007. The pain self-efficacy questionnaire: Taking pain into account. Eur J Pain, 11, 153–63.

37 Harden, N. R., Bruehl, S., Perez, R., Birklein, F., Marinus, J., Maihofner, C., Lubenow, T., Buvanendran, A., Mackey, S., Graciosa, J., Mogilevski, M., Ramsden, C., Schlereth, T., Chont, M. & Vatine, J. J. 2010. Development of a severity score for CRPS. Pain, 151, 870–876.

38 Lovibond, S.H. & Lovibond, P.F. 1995. Manual for the Depression Anxiety Stress Scales, 2nd Ed. Sydney, Australia: Psychology Foundation of Australia.

39 Sullivan, M. J. L. 1995-2009. The Pain Catastrophizing Scale - User Manual. Montreal, Quebec: McGill University

40 Neblett, R., Hartzell, M. M., Mayer, T. G., Bradford, E. M. & Gatchel, R. J. 2016. Establishing clinically meaningful severity levels for the Tampa Scale for Kinesiophobia (TSK-13). Eur J Pain, 20, 701–10.

41 O’brien, J. A., Mcguire, H. M., Shinko, D., FAZEKAS DE ST Groth, B., Russo, M. A., Bailey, D., Santarelli, D. M., Wynne, K. & Austin, P. J. 2021. T lymphocyte and monocyte subsets are dysregulated in type 1 diabetes patients with peripheral neuropathic pain. Brain Behav Immun Health, 15, 100283.

42 O’brien, J. A., Heng, B., Ahn, S. B., Staats-Pires, A., Wake, A., Wong, R., Peek, A. L., Meylakh, N., Ng, K., Stark, R., Macefield, V. G., Mcguire, H. M., Henderson, L. A. & Austin, P. J. 2025. Cellular and molecular characterisation of the peripheral immune environment in migraine. Pain Rep, 10, e1317.

43 Keyes, T. J., Koladiya, A., Lo, Y. C., Nolan, G. P. & Davis, K. L. 2023. tidytof: a user-friendly framework for scalable and reproducible high-dimensional cytometry data analysis. Bioinform Adv, 3, vbad071.

44 Olsen, L. R., Leipold, M. D., Pedersen, C. B. & Maecker, H. T. 2019. The anatomy of single cell mass cytometry data. Cytometry A, 95, 156–172.

45 Pascovici, D., Handler, D. C., Wu, J. X. & Haynes, P. A. 2016. Multiple testing corrections in quantitative proteomics: A useful but blunt tool. Proteomics, 16, 2448–53.

46 Wu, J. X., Song, X., Pascovici, D., Zaw, T., Care, N., Krisp, C. & Molloy, M. P. 2016. SWATH Mass Spectrometry Performance Using Extended Peptide MS/MS Assay Libraries. Mol Cell Proteomics, 15, 2501–14.

47 Ashburner, M., Ball, C. A., Blake, J. A., Botstein, D., Butler, H., Cherry, J. M., Davis, A. P., Dolinski, K., Dwight, S. S., Eppig, J. T., Harris, M. A., Hill, D. P., Issel-Tarver, L., Kasarskis, A., Lewis, S., Matese, J. C., Richardson, J. E., Ringwald, M., Rubin, G. M. & Sherlock, G. 2000. Gene ontology: tool for the unification of biology. The Gene Ontology Consortium. Nat Genet, 25, 25–9.

48 Pletscher-Frankild, S., Palleja, A., Tsafou, K., Binder, J. X. & Jensen, L. J. 2015. DISEASES: text mining and data integration of disease-gene associations. Methods, 74, 83–9.

49 Joshi-Tope, G., Gillespie, M., Vastrik, I., D’eustachio, P., Schmidt, E., DE Bono, B., Jassal, B., Gopinath, G. R., Wu, G. R., Matthews, L., Lewis, S., Birney, E. & Stein, L. 2005. Reactome: a knowledgebase of biological pathways. Nucleic Acids Res, 33, D428–32.

50 Kuleshov, M. V., Jones, M. R., Rouillard, A. D., Fernandez, N. F., Duan, Q., Wang, Z., Koplev, S., Jenkins, S. L., Jagodnik, K. M., Lachmann, A., Mcdermott, M. G., Monteiro, C. D., Gundersen, G. W. & MA’ayan, A. 2016. Enrichr: a comprehensive gene set enrichment analysis web server 2016 update. Nucleic Acids Res, 44, W90–7.

51 Sumera, A., Anuar, N. D., Radhakrishnan, A. K., Ibrahim, H., Rutt, N. H., Ismail, N. H., Tan, T. M. & Baba, A. A. 2020. A Novel Method to Identify Autoantibodies against Putative Target Proteins in Serum from beta-Thalassemia Major: A Pilot Study. Biomedicines, 8, 97.

52 Weber, L. M., Nowicka, M., Soneson, C. & Robinson, M. D. 2019. diffcyt: Differential discovery in high-dimensional cytometry via high-resolution clustering. Commun Biol, 2, 183.

53 DE Mos, M., DE Bruijn, A. G., Huygen, F. J., Dieleman, J. P., Stricker, B. H. & Sturkenboom, M. C. 2007. The incidence of complex regional pain syndrome: a population-based study. Pain, 129, 12–20.

54 Sandroni, P., Benrud-Larson, L. M., Mcclelland, R. L. & Low, P. A. 2003. Complex regional pain syndrome type I: incidence and prevalence in Olmsted county, a population-based study. Pain, 103, 199–207.

55 Jenks, S. A., Cashman, K. S., Zumaquero, E., Marigorta, U. M., Patel, A. V., Wang, X., Tomar, D., Woodruff, M. C., Simon, Z., Bugrovsky, R., Blalock, E. L., Scharer, C. D., Tipton, C. M., Wei, C., Lim, S. S., Petri, M., Niewold, T. B., Anolik, J. H., Gibson, G., Lee, F. E., Boss, J. M., Lund, F. E. & Sanz, I. 2018. Distinct Effector B Cells Induced by Unregulated Toll-like Receptor 7 Contribute to Pathogenic Responses in Systemic Lupus Erythematosus. Immunity, 49, 725–739 e6.

56 Sanz, I., Wei, C., Jenks, S. A., Cashman, K. S., Tipton, C., Woodruff, M. C., Hom, J. & Lee, F. E. 2019. Challenges and Opportunities for Consistent Classification of Human B Cell and Plasma Cell Populations. Front Immunol, 10, 2458.

57 Moir, S., Ho, J., Malaspina, A., Wang, W., Dipoto, A. C., O’shea, M. A., Roby, G., Kottilil, S., Arthos, J., Proschan, M. A., Chun, T. W. & Fauci, A. S. 2008. Evidence for HIV-associated B cell exhaustion in a dysfunctional memory B cell compartment in HIV-infected viremic individuals. J Exp Med, 205, 1797–805.

58 Sanz, I. 2025. Human Atypical B Cells. An Overview. Immunol Rev, 334, e70058.

59 Souto-Carneiro, M. M., Mahadevan, V., Takada, K., Fritsch-Stork, R., Nanki, T., Brown, M., Fleisher, T. A., Wilson, M., Goldbach-Mansky, R. & Lipsky, P. E. 2009. Alterations in peripheral blood memory B cells in patients with active rheumatoid arthritis are dependent on the action of tumour necrosis factor. Arthritis Res Ther, 11, R84.

60 Adlowitz, D. G., Barnard, J., Biear, J. N., Cistrone, C., Owen, T., Wang, W., Palanichamy, A., Ezealah, E., Campbell, D., Wei, C., Looney, R. J., Sanz, I. & Anolik, J. H. 2015. Expansion of Activated Peripheral Blood Memory B Cells in Rheumatoid Arthritis, Impact of B Cell Depletion Therapy, and Biomarkers of Response. PLoS One, 10, e0128269.

61 Horisberger, A., Humbel, M., Fluder, N., Bellanger, F., Fenwick, C., Ribi, C. & Comte, D. 2022. Measurement of circulating CD21(-)CD27(-) B lymphocytes in SLE patients is associated with disease activity independently of conventional serological biomarkers. Sci Rep, 12, 9189.

62 Wehr, C., Eibel, H., Masilamani, M., Illges, H., Schlesier, M., Peter, H. H. & Warnatz, K. 2004. A new CD21low B cell population in the peripheral blood of patients with SLE. Clin Immunol, 113, 161–71.

63 Jacobi, A. M., Reiter, K., Mackay, M., Aranow, C., Hiepe, F., Radbruch, A., Hansen, A., Burmester, G. R., Diamond, B., Lipsky, P. E. & Dorner, T. 2008. Activated memory B cell subsets correlate with disease activity in systemic lupus erythematosus: delineation by expression of CD27, IgD, and CD95. Arthritis Rheum, 58, 1762–73.

64 Good, K. L., Avery, D. T. & Tangye, S. G. 2009. Resting human memory B cells are intrinsically programmed for enhanced survival and responsiveness to diverse stimuli compared to naive B cells. J Immunol, 182, 890–901.

65 Sanz, I., Wei, C., Lee, F. E. & Anolik, J. 2008. Phenotypic and functional heterogeneity of human memory B cells. Semin Immunol, 20, 67–82.

66 Manz, R. A., Moser, K., Burmester, G. R., Radbruch, A. & Hiepe, F. 2006. Immunological memory stabilizing autoreactivity. Curr Top Microbiol Immunol, 305, 241–57.

67 Jenks, S. A., Cashman, K. S., Woodruff, M. C., Lee, F. E. & Sanz, I. 2019. Extrafollicular responses in humans and SLE. Immunol Rev, 288, 136–148.

68 Cowan, G. J. M., Miles, K., Capitani, L., Giguere, S. S. B., Johnsson, H., Goodyear, C., Mcinnes, I. B., Breusch, S., Gray, D. & Gray, M. 2020. In Human Autoimmunity, a Substantial Component of the B Cell Repertoire Consists of Polyclonal, Barely Mutated IgG(+ve) B Cells. Front Immunol, 11, 395.

69 Maclennan, I. C., Toellner, K. M., Cunningham, A. F., Serre, K., Sze, D. M., Zuniga, E., Cook, M. C. & Vinuesa, C. G. 2003. Extrafollicular antibody responses. Immunol Rev, 194, 8–18.

70 Li, Y., Li, Z. & Hu, F. 2021. Double-negative (DN) B cells: an under-recognized effector memory B cell subset in autoimmunity. Clin Exp Immunol, 205, 119–127.

71 Ku, S. W., Fu, T. H., You, H. L., Su, Y. J. & Huang, W. T. 2025. Alternation in Peripheral B Cell Subpopulations Is a Potential Biomarker for Autoimmune Diseases-A Cross-Sectional Study. Diagnostics (Basel), 15.

72 Wei, C., Anolik, J., Cappione, A., Zheng, B., Pugh-Bernard, A., Brooks, J., Lee, E. H., Milner, E. C. & Sanz, I. 2007. A new population of cells lacking expression of CD27 represents a notable component of the B cell memory compartment in systemic lupus erythematosus. J Immunol, 178, 6624–33.

73 Grimsholm, O. 2023. CD27 on human memory B cells-more than just a surface marker. Clin Exp Immunol, 213, 164–172.

74 Klein, U., Rajewsky, K. & Kuppers, R. 1998. Human immunoglobulin (Ig)M+IgD+ peripheral blood B cells expressing the CD27 cell surface antigen carry somatically mutated variable region genes: CD27 as a general marker for somatically mutated (memory) B cells. J Exp Med, 188, 1679–89.

75 William, J., Euler, C., Christensen, S. & Shlomchik, M. J. 2002. Evolution of autoantibody responses via somatic hypermutation outside of germinal centers. Science, 297, 2066–70.

76 Weiss, G. E., Crompton, P. D., Li, S., Walsh, L. A., Moir, S., Traore, B., Kayentao, K., Ongoiba, A., Doumbo, O. K. & Pierce, S. K. 2009. Atypical memory B cells are greatly expanded in individuals living in a malaria-endemic area. J Immunol, 183, 2176–82.

77 Bruno, T. C., Ebner, P. J., Moore, B. L., Squalls, O. G., Waugh, K. A., Eruslanov, E. B., Singhal, S., Mitchell, J. D., Franklin, W. A., Merrick, D. T., Mccarter, M. D., Palmer, B. E., Kern, J. A. & Slansky, J. E. 2017. Antigen-Presenting Intratumoral B Cells Affect CD4(+) TIL Phenotypes in Non-Small Cell Lung Cancer Patients. Cancer Immunol Res, 5, 898–907.

78 Zhu, L., Wong, Y. H., Wong, S. S. H., Cheung, S. C. Y., Sher, J. K. H., Yam, I. Y. L., Yung, S., Chan, T. M. & Yap, D. Y. H. 2024. Alterations in exhausted and classical memory B cells in lupus nephritis - Relationship with disease relapse. Clin Immunol, 265, 110284.

79 Yeo, L., Lom, H., Juarez, M., Snow, M., Buckley, C. D., Filer, A., Raza, K. & Scheel-Toellner, D. 2015. Expression of FcRL4 defines a pro-inflammatory, RANKL-producing B cell subset in rheumatoid arthritis. Ann Rheum Dis, 74, 928–35.

80 Amara, K., Clay, E., Yeo, L., Ramskold, D., Spengler, J., Sippl, N., Cameron, J. A., Israelsson, L., Titcombe, P. J., Gronwall, C., Sahbudin, I., Filer, A., Raza, K., Malmstrom, V. & Scheel-Toellner, D. 2017. B cells expressing the IgA receptor FcRL4 participate in the autoimmune response in patients with rheumatoid arthritis. J Autoimmun, 81, 34–43.

81 Wing, E., Sutherland, C., Miles, K., Gray, D., Goodyear, C. S., Otto, T. D., Breusch, S., Cowan, G. & Gray, M. 2023. Double-negative-2 B cells are the major synovial plasma cell precursor in rheumatoid arthritis. Front Immunol, 14, 1241474.

82 Sciaranghella, G., Tong, N., Mahan, A. E., Suscovich, T. J. & Alter, G. 2013. Decoupling activation and exhaustion of B cells in spontaneous controllers of HIV infection. AIDS, 27, 175–80.

83 Kardava, L., Moir, S., Wang, W., Ho, J., Buckner, C. M., Posada, J. G., O’shea, M. A., Roby, G., Chen, J., Sohn, H. W., Chun, T. W., Pierce, S. K. & Fauci, A. S. 2011. Attenuation of HIV-associated human B cell exhaustion by siRNA downregulation of inhibitory receptors. J Clin Invest, 121, 2614–24.

84 Koncz, G. & Hueber, A. O. 2012. The Fas/CD95 Receptor Regulates the Death of Autoreactive B Cells and the Selection of Antigen-Specific B Cells. Front Immunol, 3, 207.

85 Shi, J., Hou, S., Fang, Q., Liu, X., Liu, X. & Qi, H. 2018. PD-1 Controls Follicular T Helper Cell Positioning and Function. Immunity, 49, 264–274 e4.

86 Jia, X. Y., Zhu, Q. Q., Wang, Y. Y., Lu, Y., Li, Z. J., Li, B. Q., Tang, J., Wang, H. T., Song, C. W., Xie, C. H. & Chen, L. J. 2019. The role and clinical significance of programmed cell death-ligand 1 expressed on CD19(+)B-cells and subsets in systemic lupus erythematosus. Clin Immunol, 198, 89–99.

87 Nitschke, L., Carsetti, R., Ocker, B., Kohler, G. & Lamers, M. C. 1997. CD22 is a negative regulator of B-cell receptor signalling. Curr Biol, 7, 133–43.

88 Clark, E. A. & Giltiay, N. V. 2018. CD22: A Regulator of Innate and Adaptive B Cell Responses and Autoimmunity. Front Immunol, 9, 2235.

89 Hatta, Y., Tsuchiya, N., Matsushita, M., Shiota, M., Hagiwara, K. & Tokunaga, K. 1999. Identification of the gene variations in human CD22. Immunogenetics, 49, 280–6.

90 Surolia, I., Pirnie, S. P., Chellappa, V., Taylor, K. N., Cariappa, A., Moya, J., Liu, H., Bell, D. W., Driscoll, D. R., Diederichs, S., Haider, K., Netravali, I., Le, S., Elia, R., Dow, E., Lee, A., Freudenberg, J., DE Jager, P. L., Chretien, Y., Varki, A., Macdonald, M. E., Gillis, T., Behrens, T. W., Bloch, D., Collier, D., Korzenik, J., Podolsky, D. K., Hafler, D., Murali, M., Sands, B., Stone, J. H., Gregersen, P. K. & Pillai, S. 2010. Functionally defective germline variants of sialic acid acetylesterase in autoimmunity. Nature, 466, 243–7.

91 O’keefe, T. L., Williams, G. T., Batista, F. D. & Neuberger, M. S. 1999. Deficiency in CD22, a B cell-specific inhibitory receptor, is sufficient to predispose to development of high affinity autoantibodies. J Exp Med, 189, 1307–13.

92 Yarkoni, Y., Fischel, R., Kat, I., Yachimovich-Cohen, N. & Eilat, D. 2006. Peripheral B cell receptor editing may promote the production of high-affinity autoantibodies in CD22-deficient mice. Eur J Immunol, 36, 2755–67.

93 Jellusova, J., Wellmann, U., Amann, K., Winkler, T. H. & Nitschke, L. 2010. CD22 x Siglec-G double-deficient mice have massively increased B1 cell numbers and develop systemic autoimmunity. J Immunol, 184, 3618–27.

94 Matsubara, N., Imamura, A., Yonemizu, T., Akatsu, C., Yang, H., Ueki, A., Watanabe, N., Abdu-Allah, H., Numoto, N., Takematsu, H., Kitazume, S., Tedder, T. F., Marth, J. D., Ito, N., Ando, H., Ishida, H., Kiso, M. & Tsubata, T. 2018. CD22-Binding Synthetic Sialosides Regulate B Lymphocyte Proliferation Through CD22 Ligand-Dependent and Independent Pathways, and Enhance Antibody Production in Mice. Front Immunol, 9, 820.

95 Potter, K. N., Li, Y., Mageed, R. A., Jefferis, R. & Capra, J. D. 1999. Molecular characterization of the VH1-specific variable region determinants recognized by anti-idiotypic monoclonal antibodies G6 and G8. Scand J Immunol, 50, 14–20.

96 Potter, K. N., Li, Y., Pascual, V., Williams, R. C., JR., Byres, L. C., Spellerberg, M., Stevenson, F. K. & Capra, J. D. 1993. Molecular characterization of a cross-reactive idiotope on human immunoglobulins utilizing the VH4-21 gene segment. J Exp Med, 178, 1419–28.

97 Avnir, Y., Prachanronarong, K. L., Zhang, Z., Hou, S., Peterson, E. C., Sui, J., Zayed, H., Kurella, V. B., Mcguire, A. T., Stamatatos, L., Hilbert, B. J., Bohn, M. F., Kowalik, T. F., Jensen, J. D., Finberg, R. W., Wang, J. P., Goodall, M., Jefferis, R., Zhu, Q., Kurt Yilmaz, N., Schiffer, C. A. & Marasco, W. A. 2017. Structural Determination of the Broadly Reactive Anti-IGHV1-69 Anti-idiotypic Antibody G6 and Its Idiotope. Cell Rep, 21, 3243–3255.

98 Sutton, B. J., Corper, A. L., Sohi, M. K., Jefferis, R., Beale, D. & Taussig, M. J. 1998. The structure of a human rheumatoid factor bound to IgG Fc. Adv Exp Med Biol, 435, 41–50.

99 Smith, G., Spellerberg, M., Boulton, F., Roelcke, D. & Stevenson, F. 1995. The immunoglobulin VH gene, VH4-21, specifically encodes autoanti-red cell antibodies against the I or i antigens. Vox Sang, 68, 231–5.

100 Thompson, K. M., Sutherland, J., Barden, G., Melamed, M. D., Randen, I., Natvig, J. B., Pascual, V., Capra, J. D. & Stevenson, F. K. 1991. Human monoclonal antibodies against blood group antigens preferentially express a VH4-21 variable region gene-associated epitope. Scand J Immunol, 34, 509–18.

101 Silberstein, L. E., George, A., Durdik, J. M. & Kipps, T. J. 1996. The V4-34 encoded anti-i autoantibodies recognize a large subset of human and mouse B-cells. Blood Cells Mol Dis, 22, 126–38.

102 Bhat, N. M., Bieber, M. M., Spellerberg, M. B., Stevenson, F. K. & Teng, N. N. 2000. Recognition of auto- and exoantigens by V4-34 gene encoded antibodies. Scand J Immunol, 51, 134–40.

103 Cappione, A. J., Pugh-Bernard, A. E., Anolik, J. H. & Sanz, I. 2004. Lupus IgG VH4.34 antibodies bind to a 220-kDa glycoform of CD45/B220 on the surface of human B lymphocytes. J Immunol, 172, 4298–307.

104 VAN Vollenhoven, R. F., Bieber, M. M., Powell, M. J., Gupta, P. K., Bhat, N. M., Richards, K. L., Albano, S. A. & Teng, N. N. 1999. VH4-34 encoded antibodies in systemic lupus erythematosus: a specific diagnostic marker that correlates with clinical disease characteristics. J Rheumatol, 26, 1727–33.

105 Bhat, N. M., Lee, L. M., VAN Vollenhoven, R. F., Teng, N. N. & Bieber, M. M. 2002. VH4-34 encoded antibody in systemic lupus erythematosus: effect of isotype. J Rheumatol, 29, 2114–21.

106 Richardson, C., Chida, A. S., Adlowitz, D., Silver, L., Fox, E., Jenks, S. A., Palmer, E., Wang, Y., Heimburg-Molinaro, J., Li, Q. Z., Mohan, C., Cummings, R., Tipton, C. & Sanz, I. 2013. Molecular basis of 9G4 B cell autoreactivity in human systemic lupus erythematosus. J Immunol, 191, 4926–39.

107 Reed, J. H., Jackson, J., Christ, D. & Goodnow, C. C. 2016. Clonal redemption of autoantibodies by somatic hypermutation away from self-reactivity during human immunization. J Exp Med, 213, 1255–65.

108 Tipton, C. M., Fucile, C. F., Darce, J., Chida, A., Ichikawa, T., Gregoretti, I., Schieferl, S., Hom, J., Jenks, S., Feldman, R. J., Mehr, R., Wei, C., Lee, F. E., Cheung, W. C., Rosenberg, A. F. & Sanz, I. 2015. Diversity, cellular origin and autoreactivity of antibody-secreting cell population expansions in acute systemic lupus erythematosus. Nat Immunol, 16, 755–65.

109 Deguine, J. & Xavier, R. J. 2024. B cell tolerance and autoimmunity: Lessons from repertoires. J Exp Med, 221.

110 Weller, S., Faili, A., Garcia, C., Braun, M. C., LE Deist, F. F., DE SAINT Basile, G. G., Hermine, O., Fischer, A., Reynaud, C. A. & Weill, J. C. 2001. CD40-CD40L independent Ig gene hypermutation suggests a second B cell diversification pathway in humans. Proc Natl Acad Sci U S A, 98, 1166–70.

111 Shi, Y., Agematsu, K., Ochs, H. D. & Sugane, K. 2003. Functional analysis of human memory B-cell subpopulations: IgD+CD27+ B cells are crucial in secondary immune response by producing high affinity IgM. Clin Immunol, 108, 128–37.

112 Ushijima, T. S., Teruya, H., Abe, T., Itamiya, T., Ota, M., Okamura, T. & Fujio, K. 2025. Stratification of systemic lupus erythematosus with IGHV4-34 in unswitched memory B cells. Rheumatology (Oxford), 64, 6099–6105.

113 Daha, N. A., Banda, N. K., Roos, A., Beurskens, F. J., Bakker, J. M., Daha, M. R. & Trouw, L. A. 2011. Complement activation by (auto-) antibodies. Mol Immunol, 48, 1656–65.

114 Shi, X., Guo, T. Z., Li, W. W., Birklein, F., Escolano, F. L., Herrnberger, M., Clark, J. D. & Kingery, W. S. 2021. C5a complement and cytokine signaling mediate the pronociceptive effects of complex regional pain syndrome patient IgM in fracture mice. Pain, 162, 1400–1415.

115 Clark, J. D., Tawfik, V. L., Tajerian, M. & Kingery, W. S. 2018. Autoinflammatory and autoimmune contributions to complex regional pain syndrome. Mol Pain, 14, 1744806918799127.

116 Butterworth, M., Mcclellan, B. & Allansmith, M. 1967. Influence of sex in immunoglobulin levels. Nature, 214, 1224–5.

117 Fairweather, D., Frisancho-Kiss, S. & Rose, N. R. 2008. Sex differences in autoimmune disease from a pathological perspective. Am J Pathol, 173, 600–9.

118 Guo, T. Z., Shi, X., Li, W. W., Wei, T., Clark, J. D. & Kingery, W. S. 2019. Sex differences in the temporal development of pronociceptive immune responses in the tibia fracture mouse model. Pain, 160, 2013–2027.

119 Lee, H. J., Remacle, A. G., Hullugundi, S. K., Dolkas, J., Leung, J. B., Chernov, A. V., Yaksh, T. L., Strongin, A. Y. & Shubayev, V. I. 2022. Sex-Specific B Cell and Anti-Myelin Autoantibody Response After Peripheral Nerve Injury. Front Cell Neurosci, 16, 835800.

120 Linher-Melville, K., Shah, A. & Singh, G. 2020. Sex differences in neuro(auto)immunity and chronic sciatic nerve pain. Biol Sex Differ, 11, 62.

121 Goebel, A., Leite, M. I., Yang, L., Deacon, R., Cendan, C. M., FOX-Lewis, A. & Vincent, A. 2011. The passive transfer of immunoglobulin G serum antibodies from patients with longstanding Complex Regional Pain Syndrome. Eur J Pain, 15, 504 e1–6.

122 Goebel, A., Krock, E., Gentry, C., Israel, M. R., Jurczak, A., Urbina, C. M., Sandor, K., Vastani, N., Maurer, M., Cuhadar, U., Sensi, S., Nomura, Y., Menezes, J., Baharpoor, A., Brieskorn, L., Sandstrom, A., Tour, J., Kadetoff, D., Haglund, L., Kosek, E., Bevan, S., Svensson, C. I. & Andersson, D. A. 2021. Passive transfer of fibromyalgia symptoms from patients to mice. J Clin Invest, 131.

123 Liu, T., Zhang, L., Joo, D. & Sun, S. C. 2017. NF-kappaB signaling in inflammation. Signal Transduct Target Ther, 2, 17023-.

124 Makarov, S. S. 2001. NF-kappa B in rheumatoid arthritis: a pivotal regulator of inflammation, hyperplasia, and tissue destruction. Arthritis Res, 3, 200–6.

125 Oh, H. & Ghosh, S. 2013. NF-kappaB: roles and regulation in different CD4(+) T-cell subsets. Immunol Rev, 252, 41–51.

126 Balasubramani, A., Shibata, Y., Crawford, G. E., Baldwin, A. S., Hatton, R. D. & Weaver, C. T. 2010. Modular utilization of distal cis-regulatory elements controls Ifng gene expression in T cells activated by distinct stimuli. Immunity, 33, 35–47.

127 Groom, J. R. & Luster, A. D. 2011. CXCR3 in T cell function. Exp Cell Res, 317, 620–31.

128 Uceyler, N., Eberle, T., Rolke, R., Birklein, F. & Sommer, C. 2007. Differential expression patterns of cytokines in complex regional pain syndrome. Pain, 132, 195–205.

129 Lenz, M., Uceyler, N., Frettloh, J., Hoffken, O., Krumova, E. K., Lissek, S., Reinersmann, A., Sommer, C., Stude, P., Waaga-Gasser, A. M., Tegenthoff, M. & Maier, C. 2013. Local cytokine changes in complex regional pain syndrome type I (CRPS I) resolve after 6 months. Pain, 154, 2142–2149.

130 Austin, P. J., Kim, C. F., Perera, C. J. & Moalem-Taylor, G. 2012. Regulatory T cells attenuate neuropathic pain following peripheral nerve injury and experimental autoimmune neuritis. Pain, 153, 1916–1931.

131 Lees, J. G., Duffy, S. S., Perera, C. J. & Moalem-Taylor, G. 2015. Depletion of Foxp3+ regulatory T cells increases severity of mechanical allodynia and significantly alters systemic cytokine levels following peripheral nerve injury. Cytokine, 71, 207–14.

132 Fiore, N. T., Debs, S. R., Hayes, J. P., Duffy, S. S. & Moalem-Taylor, G. 2023. Pain-resolving immune mechanisms in neuropathic pain. Nat Rev Neurol, 19, 199–220.

133 Mold, C., Gewurz, H. & DU Clos, T. W. 1999. Regulation of complement activation by C-reactive protein. Immunopharmacology, 42, 23–30.

134 Lu, J., Marnell, L. L., Marjon, K. D., Mold, C., DU Clos, T. W. & Sun, P. D. 2008. Structural recognition and functional activation of FcgammaR by innate pentraxins. Nature, 456, 989–92.

135 Matsushita, M. 2013. Ficolins in complement activation. Mol Immunol, 55, 22–6.

136 Sproston, N. R. & Ashworth, J. J. 2018. Role of C-Reactive Protein at Sites of Inflammation and Infection. Front Immunol, 9, 754.

137 Pope, J. E. & Choy, E. H. 2021. C-reactive protein and implications in rheumatoid arthritis and associated comorbidities. Semin Arthritis Rheum, 51, 219–229.

138 Lindelof, L., Rantapaa-Dahlqvist, S., Lundtoft, C., Sandling, J. K., Leonard, D., Sayadi, A., Ronnblom, L., Enocsson, H., Sjowall, C., Jonsen, A., Bengtsson, A. A., Hong, M. G., Diaz-Gallo, L. M., Bianchi, M., Kozyrev, S. V., Lindblad-Toh, K., Consortium, D., Immunoarray, C., NILSSON Ekdahl, K., Nilsson, B., Gunnarsson, I., Svenungsson, E. & Eriksson, O. 2024. A survey of ficolin-3 activity in Systemic Lupus Erythematosus reveals a link to hematological disease manifestations and autoantibody profile. J Autoimmun, 143, 103166.

139 Guo, T. Z., Wei, T., Shi, X., Yuan, H., Goebel, A., Neiland, H., Birklein, F., Rebhorn, C., Clark, J. D., Li, W. W., & Kingery, W. S. 2026. Chronic complex regional pain syndrome patient IgM has pronociceptive effects in the spinal cord, but not skin, of fracture mice. [preprint]. 10.21203/rs.3.rs-11099287/v1.

140 Bergdolt, S., Kovtun, A., Hagele, Y., Liedert, A., Schinke, T., Amling, M., Huber-Lang, M. & Ignatius, A. 2017. Osteoblast-specific overexpression of complement receptor C5aR1 impairs fracture healing. PLoS One, 12, e0179512.

141 Norin, U., Rintisch, C., Meng, L., Forster, F., Ekman, D., Tuncel, J., Klocke, K., Backlund, J., Yang, M., Bonner, M. Y., Lahore, G. F., James, J., Shchetynsky, K., Bergquist, M., Gjertsson, I., Hubner, N., Backdahl, L. & Holmdahl, R. 2021. Endophilin A2 deficiency protects rodents from autoimmune arthritis by modulating T cell activation. Nat Commun, 12, 610.

142 Rice, M. R., Matta, B., Wang, L., Luo, Q., DE Guzman, J., Srinivasan, D., Ludwig, K. R., Indukuri, S., Brune, L., Tan, S. L. & Barnes, B. J. 2025. TLR-induced STK25 activation promotes IRF5-mediated inflammation. Life Sci Alliance, 8.

143 Sieghart, D., Konrad, C., Swiniarski, S., Haslacher, H., Aletaha, D. & Steiner, G. 2022. The diagnostic and prognostic value of IgG and IgA anti-citrullinated protein antibodies in patients with early rheumatoid arthritis. Front Immunol, 13, 1096866.

144 Svard, A., Kastbom, A., Soderlin, M. K., Reckner-Olsson, A. & Skogh, T. 2011. A comparison between IgG- and IgA-class antibodies to cyclic citrullinated peptides and to modified citrullinated vimentin in early rheumatoid arthritis and very early arthritis. J Rheumatol, 38, 1265–72.

