## Supplementary materials for "B cell tolerance is dysregulated in people with complex regional pain syndrome: A high-dimensional immunophenotyping study"

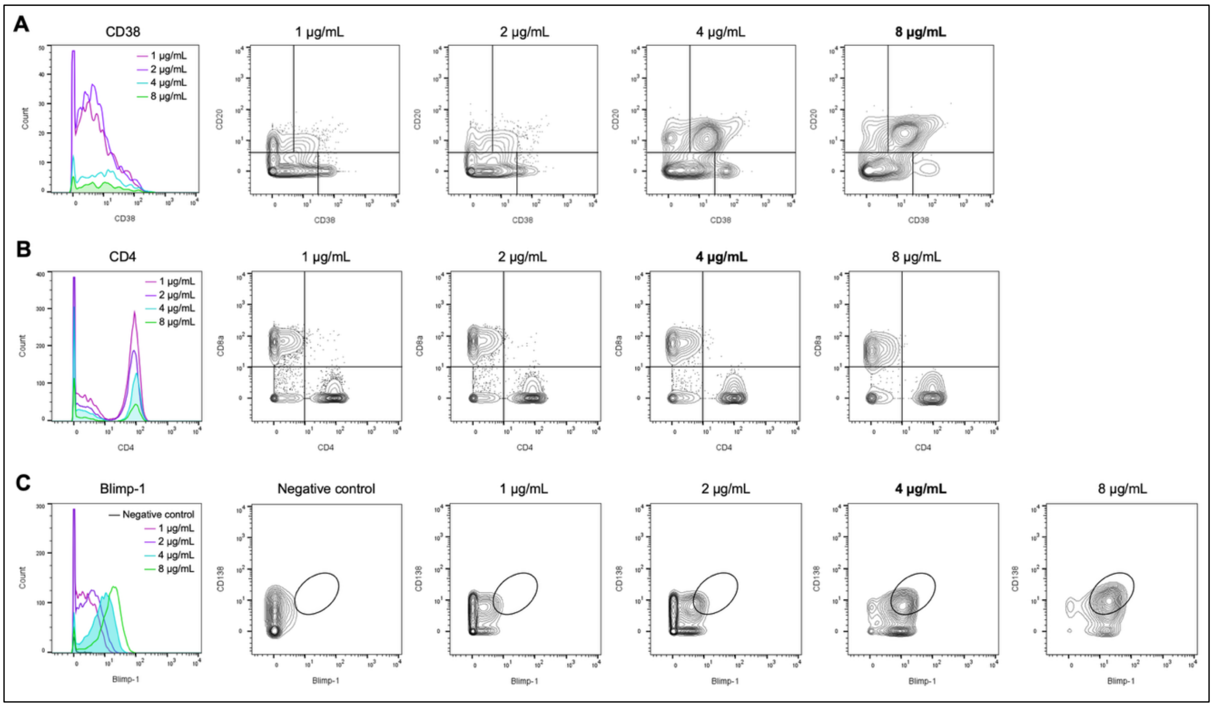


**Supplementary Figure 1.** Representative optimisation strategy for select CyTOF antibodies used to assess major peripheral B cell and T cell subsets. Healthy donor PBMCs were stained with 1, 2, 4, or 8 μg/mL of each antibody and acquired on a Helios mass cytometer. A negative control sample (0 μg/mL) was also included for intracellular and nuclear markers (such as Blimp-1; panel **C**) to confirm targeted binding. Each sample was then analysed by manual gating in FlowJo to determine the staining concentration that produced the most distinct cell populations. FlowJo histograms and countour plots show the expression of different markers at each test concentration. **(A)** Surface CD38 expression was analysed within the CD19$^{+}$CD3$^{-}$ B cell gate, **(B)** surface CD4 expression within the CD3$^{+}$CD19$^{-}$ T cell gate, and **(C)** intracellular Blimp-1 staining within the CD19$^{+}$CD3$^{-}$ B cell gate. Shaded histogram areas and bold text represent the optimal staining concentration selected for each antibody.

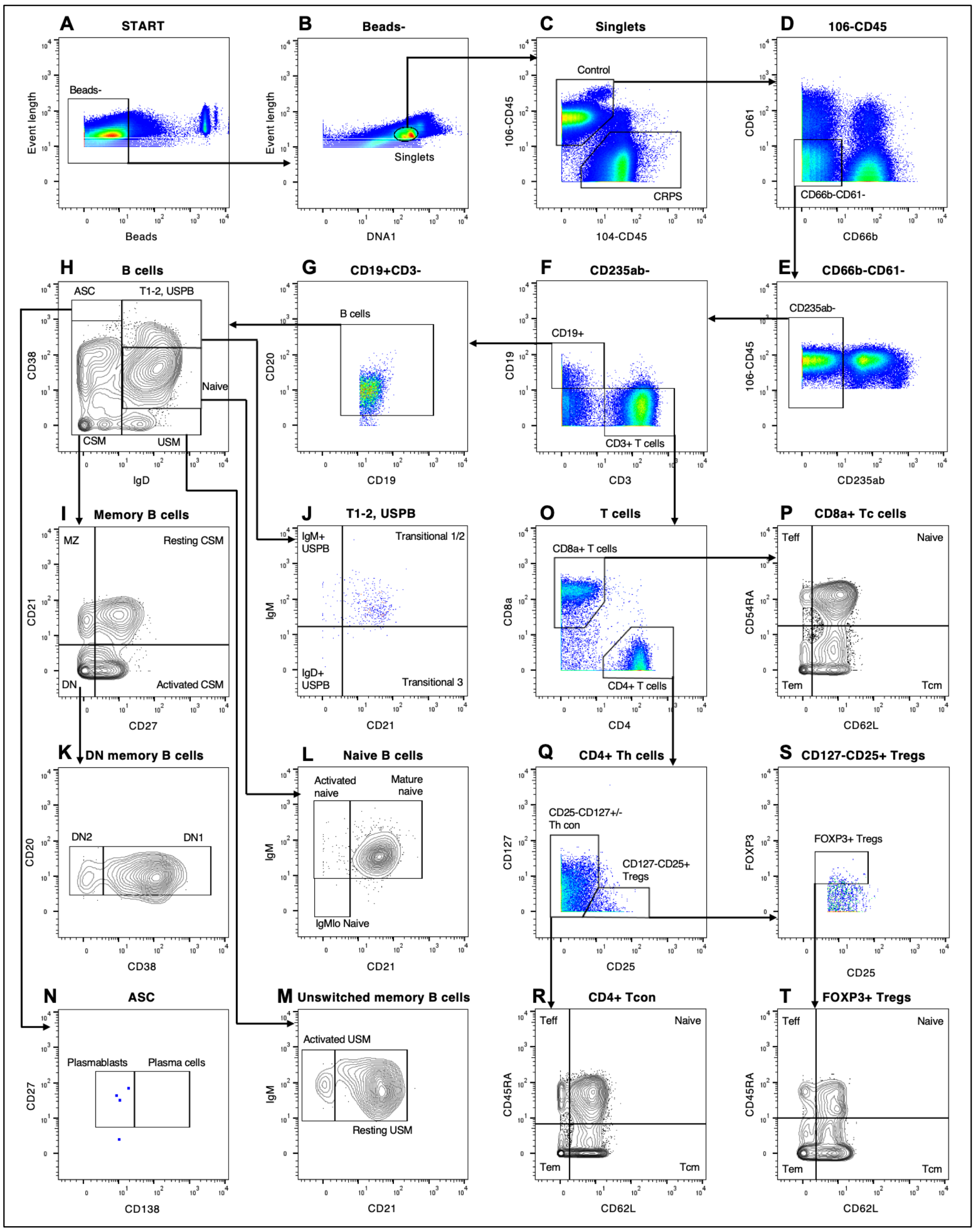


**Supplementary Figure 2.** Manual gating strategy used to identify major peripheral B cell and T cell subsets in mass cytometry experiments. For each sample, **(A)** normalisation beads were first excluded, then **(B)** cell singlets were selected, and **(C)** CRPS and control samples de-barcoded based on 104-CD45 and 106-CD45 expression, respectively. **(D-E)** Platelets, granulocytes, and erythrocytes were excluded, then **(F)** T and B lymphocyte compartments were isolated using core lineage markers ($CD3^{+}CD19^{-}$ for T cells; $CD3^{-}CD19^{+}CD20^{+}$ for B cells). **(G-T)** B cells and T cells were gated into known subsets according to core surface marker expression. Lymphocyte population statistics were then exported for frequency analysis. T cell gates downstream of CD4/CD8a **(P-T)** were included only in the first mass cytometry experiment. All gates were set manually using pseudocolour and contour plots (shown) in FlowJo (v10.10.0, BD). Gates shown are from one representative control sample. Boxes and arrows indicate cell populations passed on for analysis in subsequent gates. ASC = antibody-secreting cell; CSM = class-switched memory; DN = double-negative; MZ = marginal zone; USM = unswitched memory; USPB = unswitched plasmablast; Tc = cytotoxic T cell; Tcm = T central memory; Teff = T effector; Tem = T effector memory; Th = T helper; Th con = conventional helper T cell; Treg = regulatory T cell; T1-2 = transitional B cells type 1-2.

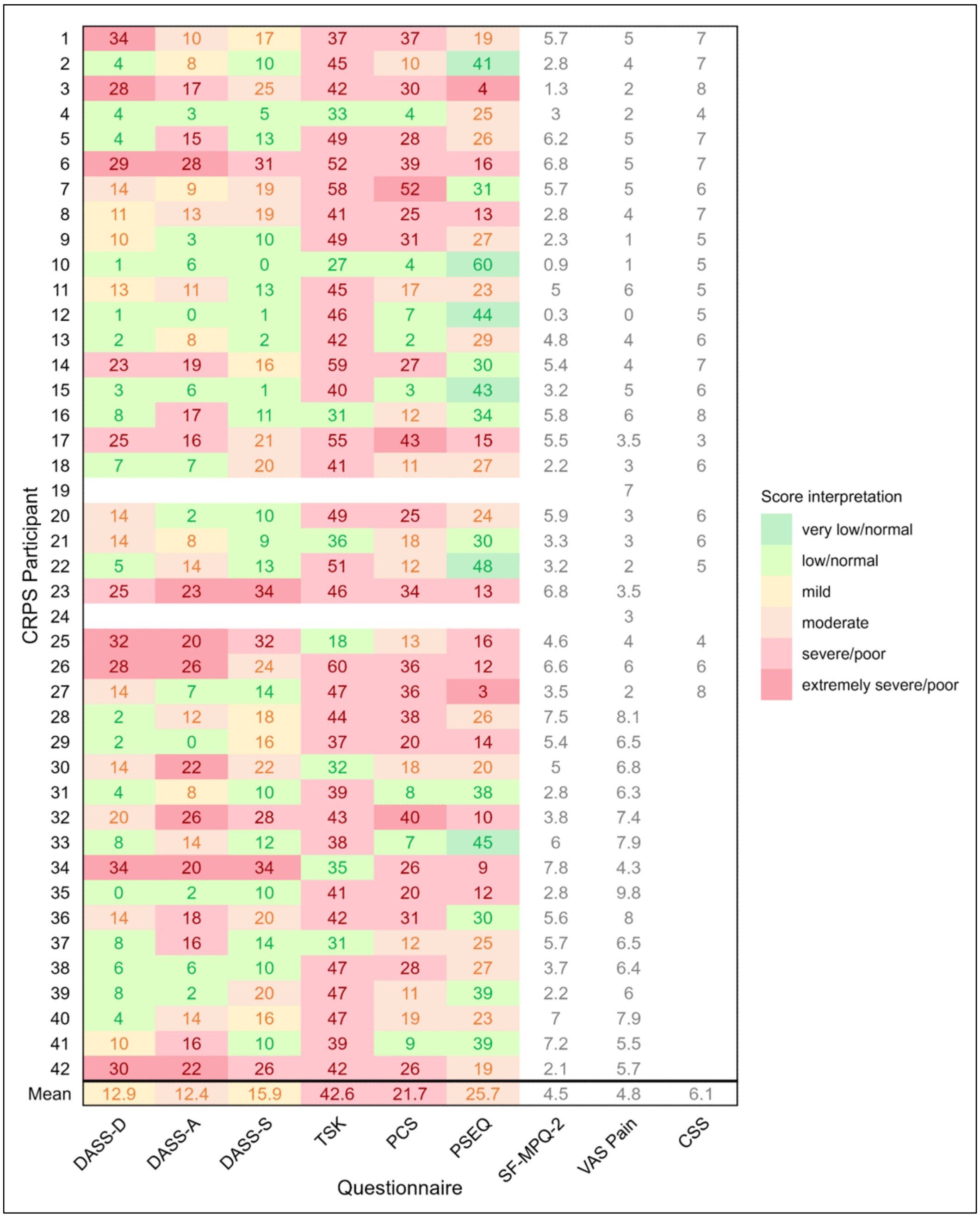


**Supplementary Figure 3.** Results of pain and psychological symptoms assessment in the CRPS cohort ($n$ = 42). The heatmap shows individual participant scores (and group mean) for each questionnaire, with colours corresponding to the score's interpretation range (e.g. mild, moderate, severe), as specified by published guidelines [1-4]. White cells are shown for measures without specific score interpretation guidelines (SF-MPQ-2 and VAS pain rating), and empty cells are shown where the participant did not provide questionnaire data. CSS = CRPS Symptoms Scale (patient-reported section only); DASS = Depression, Anxiety, Stress Scale; PCS = Pain Catastrophising Scale; PSEQ = Pain Self-Efficacy Questionnaire; SF-MPQ-2 = revised Short-Form McGill Pain Questionnaire; TSK = Tampa Scale for Kinesiophobia; VAS = Visual-Analogue Scale.

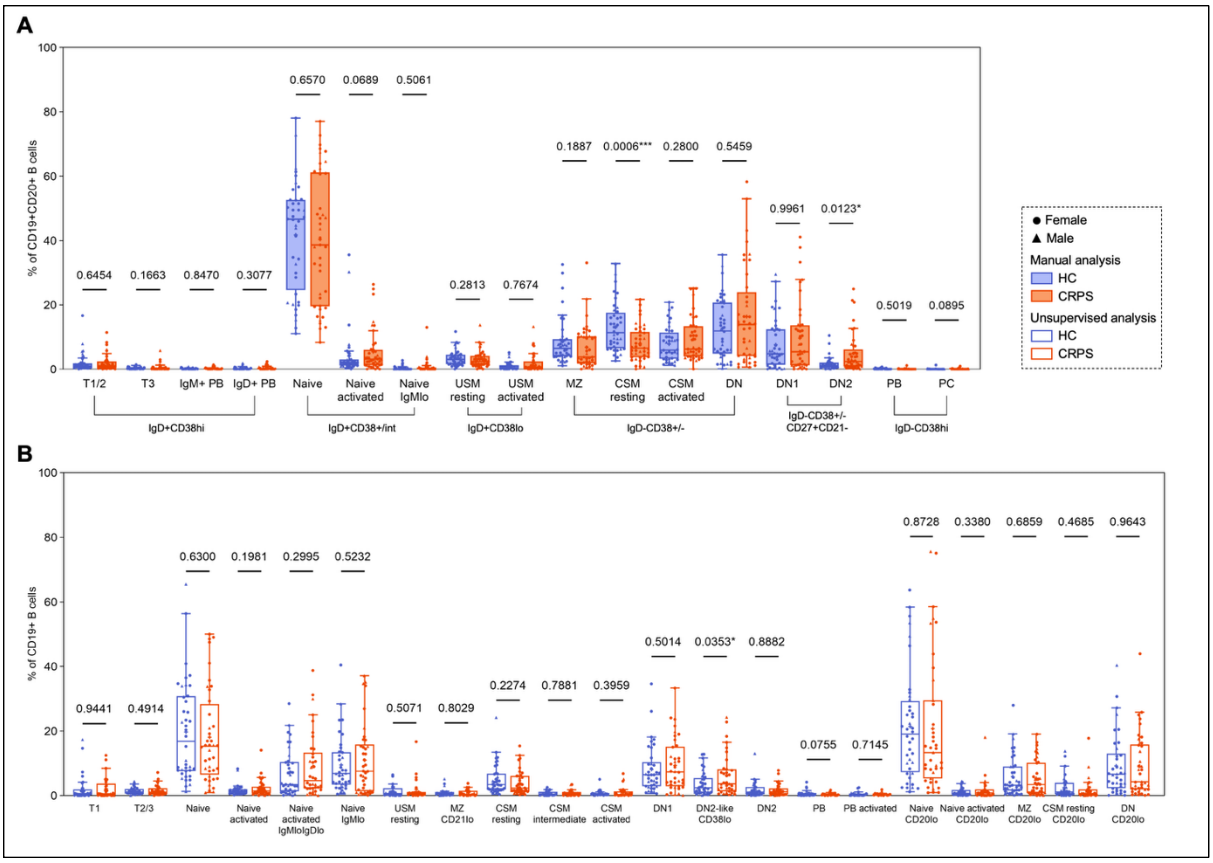


**Supplementary Figure 4.** Results of peripheral B cell profiling between CRPS subjects ($n$ = 41) and healthy donors ($n$ = 40) in the follow-up mass cytometry study. Boxplots show IQR, median (solid lines), and mean (+), with whiskers extending to 1.5x IQR, for **(A)** manually gated B cell subsets and **(B)** $CD19^+$ FlowSOM metaclusters. Statistical comparisons were performed using Welch's t-tests for parametric data (determined by Shapiro-Wilk test) or Mann-Whitney U-tests for non-parametric data, with *P < 0.05; ***P < 0.001 (Holm-corrected P values shown for unsupervised analysis). Statistical outliers in manual data were excluded using ROUT (Q = 1%). Each point represents an individual participant. CSM = class-switched memory; DN1/2 = double-negative type 1, 2; MZ = marginal zone; PB = plasmablast; PC = plasma cell; T1/2/3 = transitional type 1, 2, 3; USM = unswitched memory.

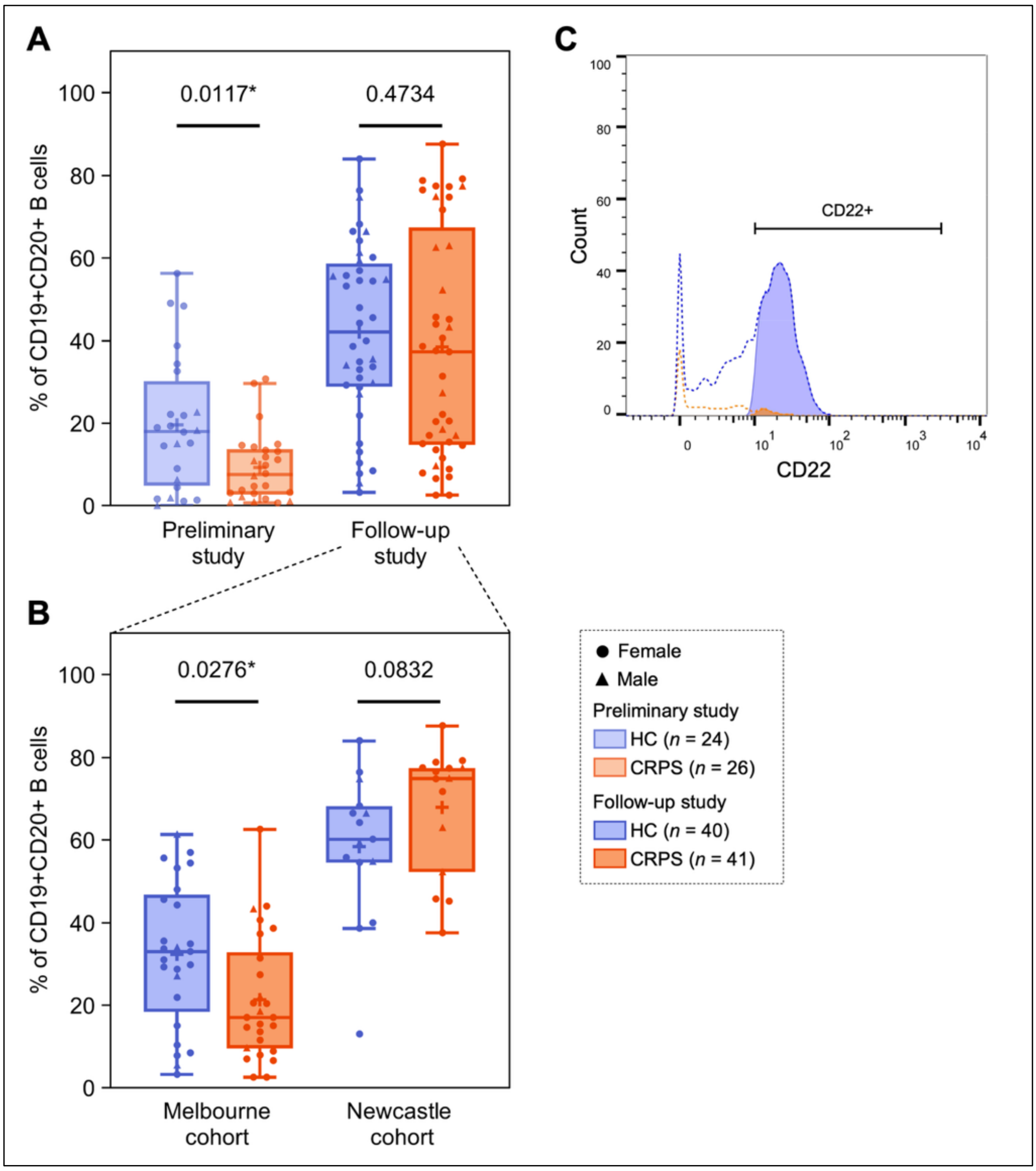


**Supplementary Figure 5.** Different proportions of B cells expressing the inhibitory co-receptor CD22 between CRPS subjects and healthy donors profiled during preliminary and follow-up mass cytometry studies. The preliminary study included samples from 27 CRPS participants ($n = 26$ acquired) collected between 2022 and 2024 at the Baker Heart and Diabetes Institute (Melbourne, NSW). The same samples from this "Melbourne cohort" were profiled in the follow-up study, along with an additional cohort of CRPS patient samples ($n = 15$) collected between 2017 and 2018 at Genesis Research Services (Broadmeadow, NSW) ("Newcastle cohort"). Reduced CD22 expression was seen in CRPS subjects from the Melbourne cohort during the **(A)** the preliminary CyTOF study and **(B)** the follow-up CyTOF study, while CRPS subjects from the Newcastle cohort (follow-up study) showed similar CD22 expression to their matched controls ($n = 15$). $CD22^+$ B cell populations were identified manually in FlowJo according to the strategy shown in panel **(C)**, then compared between CRPS and controls as a proportion of the total B cell compartment. Statistical comparisons were performed using Welch's t-tests for parametric data (as determined by Shapiro-Wilk test) or Mann-Whitney U-tests for non-parametric data, with $*P < 0.05$. Statistical outliers were excluded using ROUT (Q = 1%). Boxplots show IQR, median (solid lines), and mean (+), with whiskers extending to 1.5x IQR. Each point represents an individual participant.

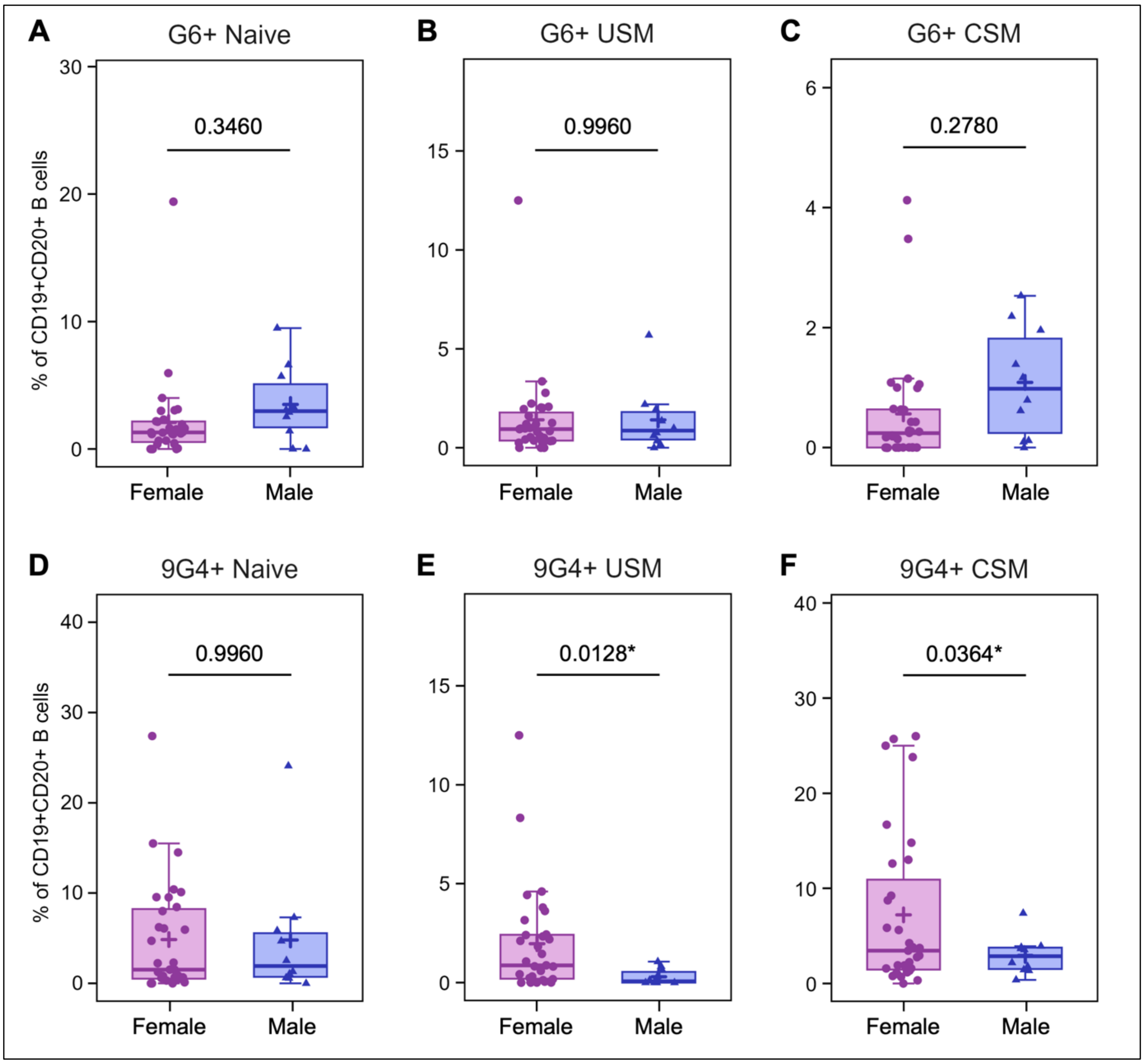


**Supplementary Figure 6.** Higher 9G4 [IGHV4-34] binding in the USM and CSM B cell compartments of women with CRPS (*n* = 31) compared to men (*n* = 11). Statistical comparisons were performed using Welch's t-tests, with *$P_{adj}$ < 0.05 (corrected for multiple comparisons using Benjamini-Hochberg method). Boxplots show IQR, median (solid lines), and mean (+) for each cell subpopulation, with whiskers extending to 1.5x IQR. Each point represents an individual CRPS participant. USM = unswitched memory; CSM = class-switched memory.

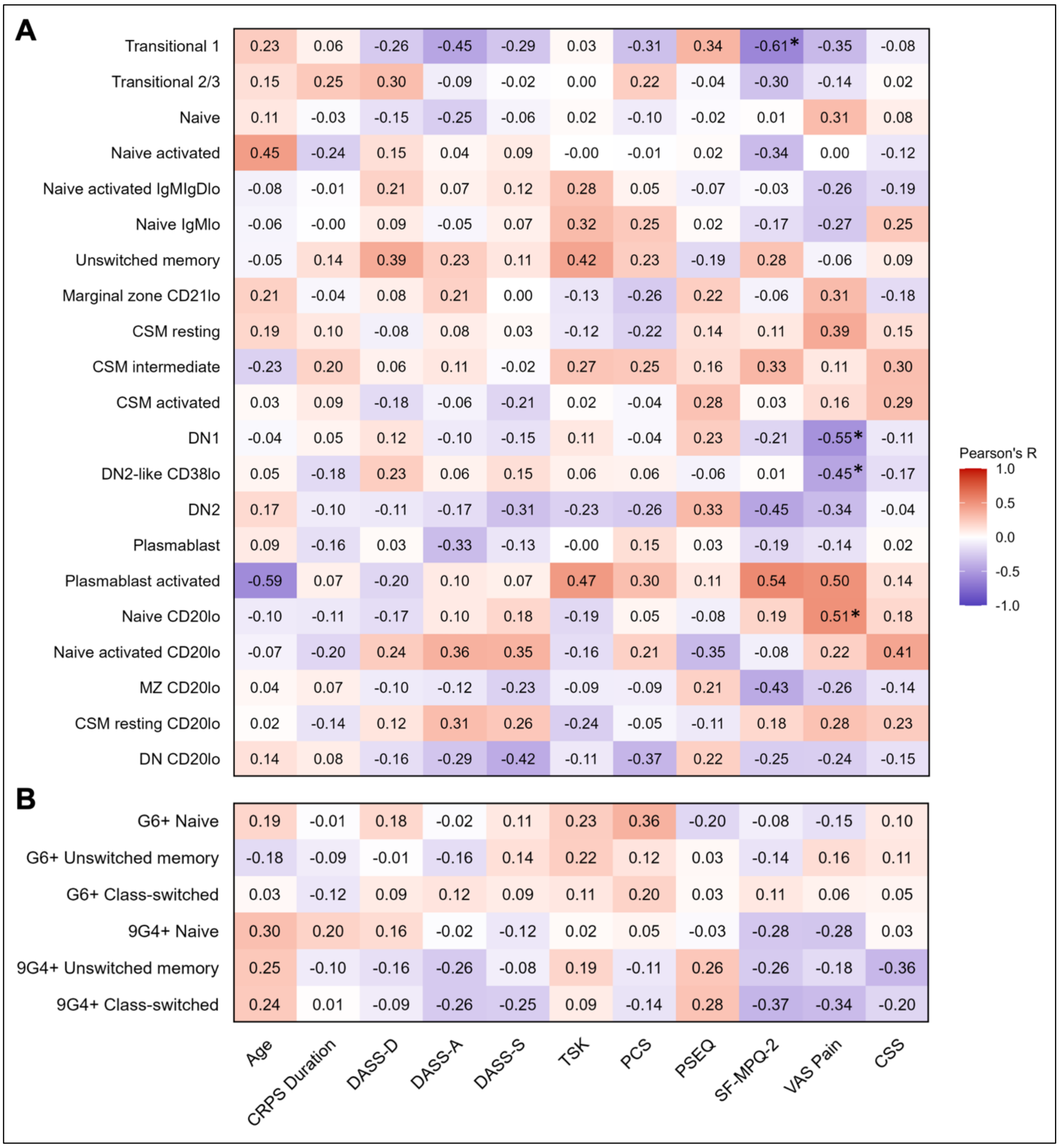


**Supplementary Figure 7.** Results of Pearson's correlation analysis between CRPS participant pain and psychological scores with major B cell subpopulations identified during the follow-up mass cytometry study. Heatmaps show correlations between clinical variables of the CRPS cohort and the proportion of **(A)** $CD19^{+}$ B cell metaclusters identified by unsupervised FlowSOM analysis, and **(B)** manually gated G6 and 9G4 B cell subpopulations. Colour represents Pearson's R value (red = positive direction, blue = negative direction), with $^{*}P_{adj} < 0.05$ (corrected for multiple comparisons using Benjamini-Hochberg method).

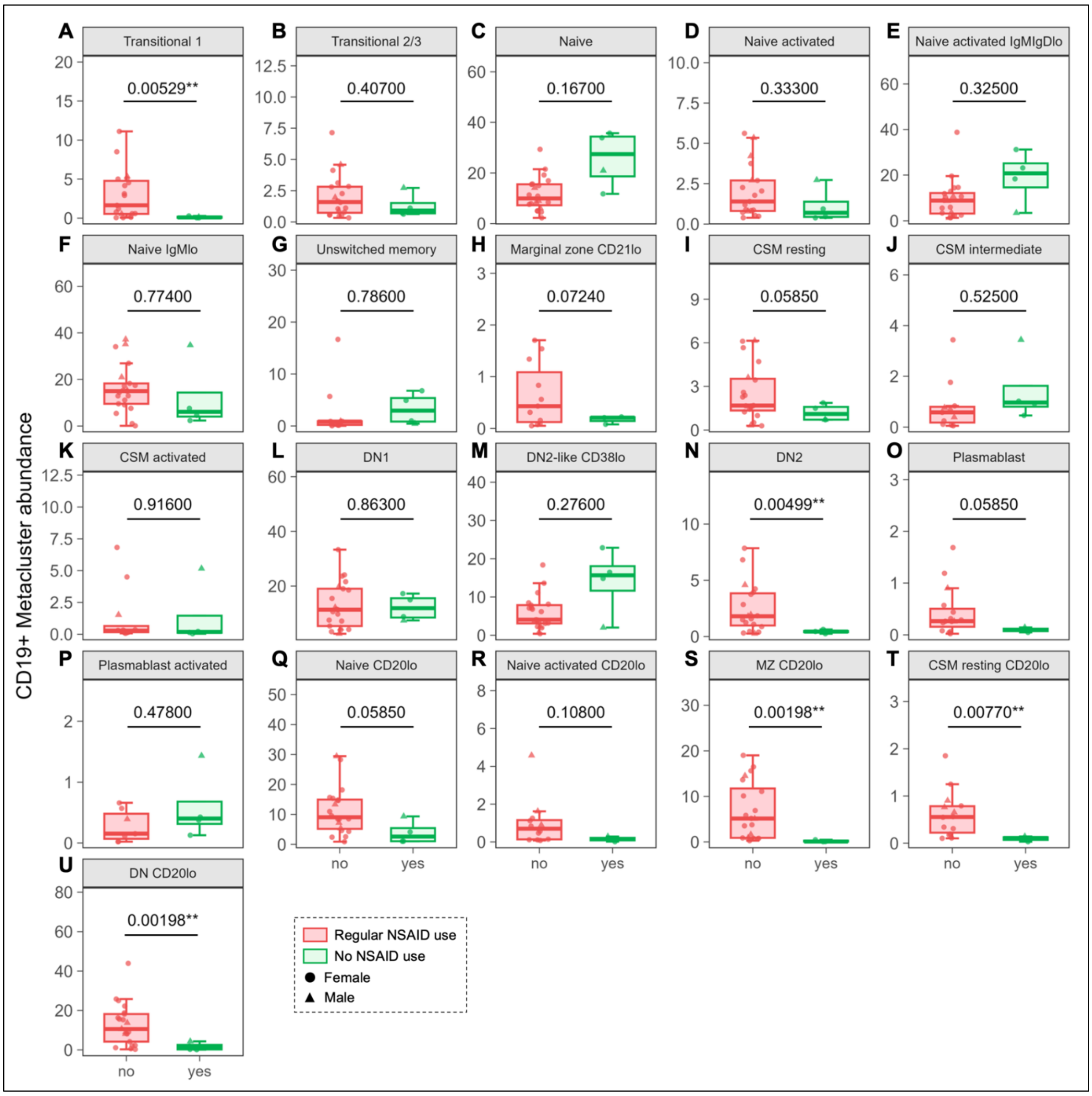


**Supplementary Figure 8.** CRPS participants reporting regular use of NSAIDs e.g. ibuprofen, naproxen celecoxib, meloxicam ($n$ = 14; $n$ = 12 completed the 7-day drug washout before blood collection) had lower proportions of transitional 1 B cells (**A**), DN2 memory B cells (**N**), as well as several CD20$^{lo}$ subsets (**S-U**) compared to CRPS participants who did not report regular NSAID use ($n$ = 12). Statistical comparisons were performed using Welch's t-tests, with $^{**}P_{adj} < 0.01$ (corrected for multiple comparisons using Benjamini-Hochberg method). Boxplots show IQR and median (bold lines), with whiskers extending to 1.5x IQR. Each point represents an individual CRPS participant.

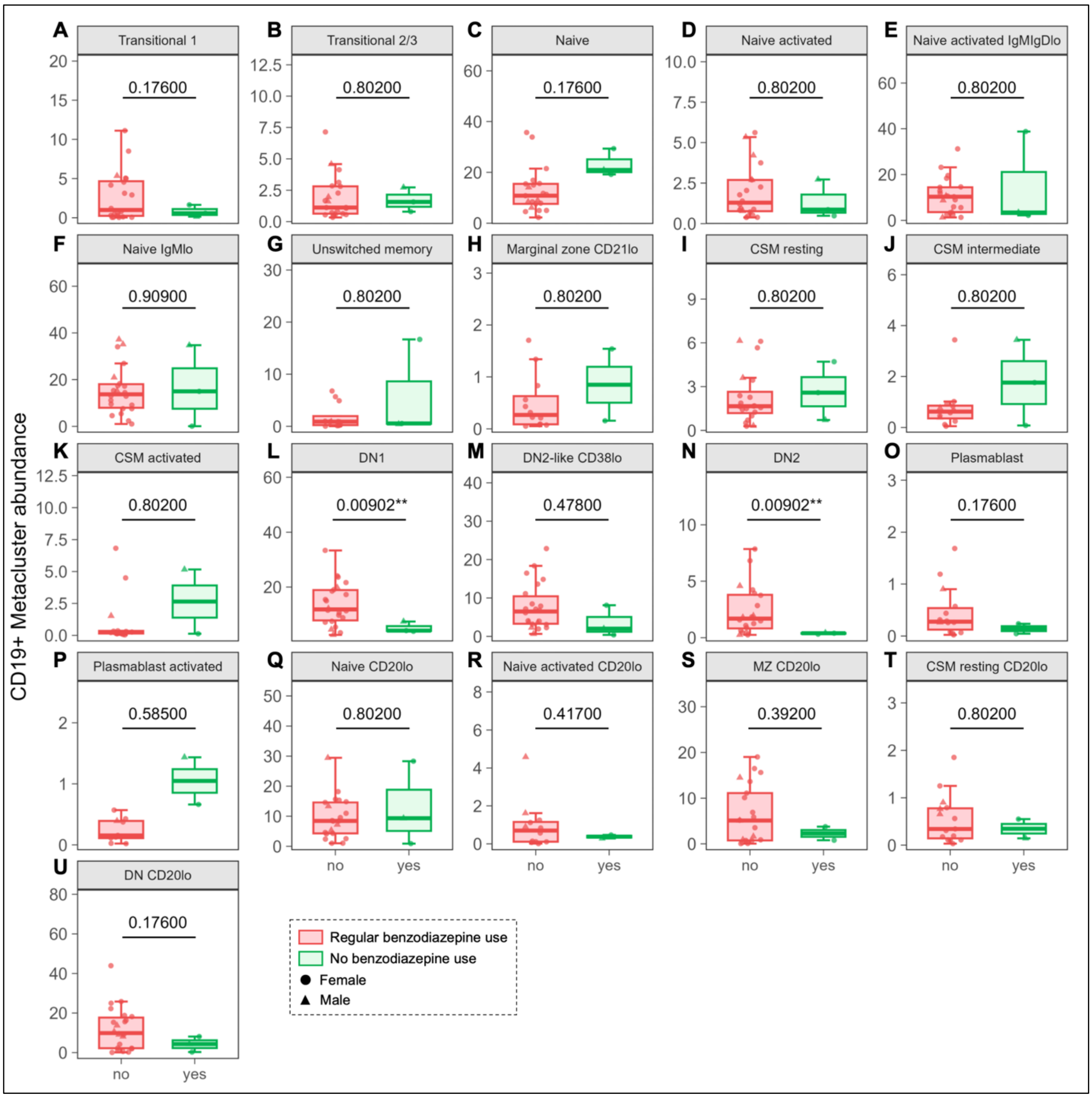


**Supplementary Figure 9.** CRPS participants reporting regular use of benzodiazepines e.g. clonazepam, diazepam (*n* = 3; all completed the 7-day drug washout before blood collection) had lower proportions of DN1 memory B cells (**L**) and DN2 memory B cells (**N**) compared to CRPS participants who did not report regular benzodiazepine use (*n* = 23). Statistical comparisons were performed using Welch's t-tests, with $**P_{adj} < 0.01$ (corrected for multiple comparisons using Benjamini-Hochberg method). Boxplots show IQR and median (bold lines), with whiskers extending to 1.5x IQR. Each point represents an individual CRPS participant.

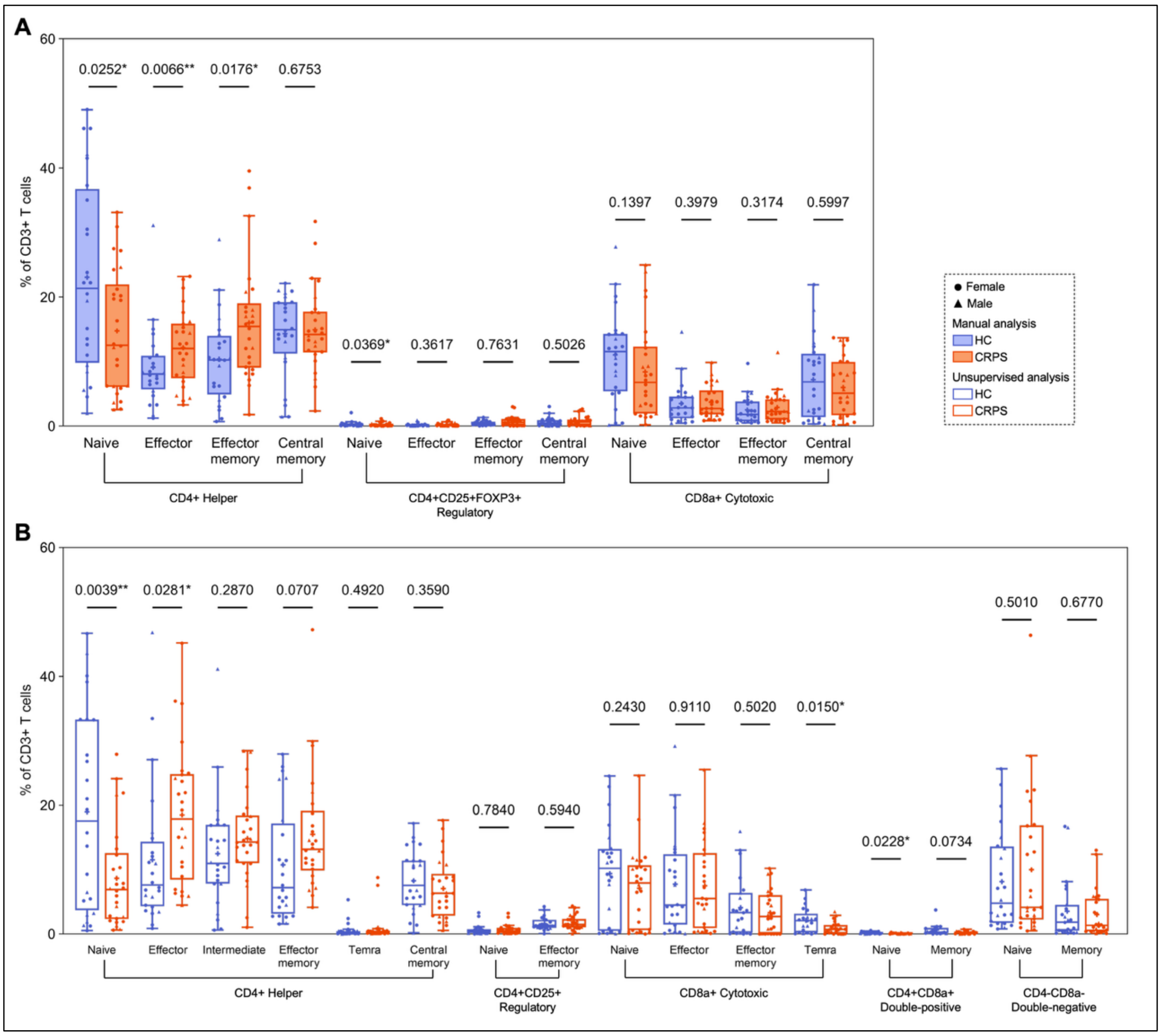


**Supplementary Figure 10.** Results of peripheral T cell profiling between CRPS subjects (*n* = 26) and healthy controls (*n* = 24) in the preliminary mass cytometry study. Boxplots show IQR, median (solid lines), and mean (+), with whiskers extending to 1.5x IQR, of **(A)** manually gated T cell subsets and **(B)** $CD3^+$ FlowSOM metaclusters, shown as a proportion of the total $CD3^+$ T cell compartment. Statistical comparisons were performed using Welch's t-tests for parametric data (determined by Shapiro-Wilk test) or Mann-Whitney U-tests for non-parametric data, with *P < 0.05; **P < 0.01 (Holm-corrected P values shown for unsupervised analysis). Statistical outliers in manual data were excluded using ROUT (Q = 1%). Each point represents an individual participant.

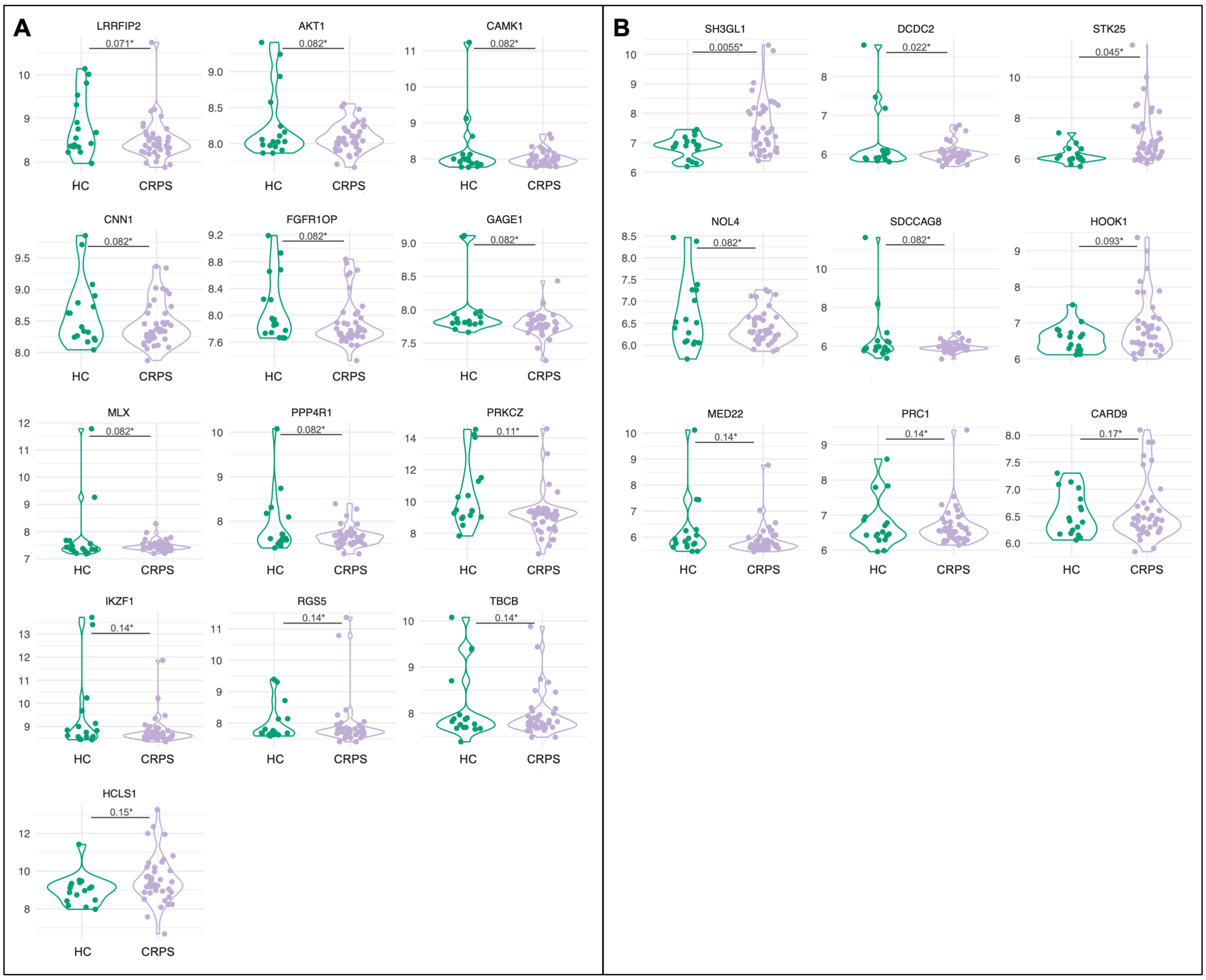


**Supplementary Figure 11.** Top antigens from the iOme® Discovery Protein Array identified by penetrance fold change analysis of serum autoantibody profiles between CRPS subjects (n = 42) and healthy donors (n = 18). Violin plots show distribution of **(A)** IgG and **(B)** IgA autoantibody levels across all samples for top antigens identified by pFC analysis of CRPS and healthy donor serum, defined as pFC ≥ 2.0 and P < 0.2 (Fisher's exact test). Each point represents an individual participant. *P < 0.2.

**Supplementary Table 1.** CyTOF antibodies used to profile major circulating B cell and T cell subsets during the preliminary mass cytometry study.

| Label | Target | Clone | Supplier | Catalogue # | Use |
|---|---|---|---|---|---|
| 89Y | IgD | IA6-2 | BD | 555776 | Phenotype |
| 104Pd | CD45 | HI30 | Biolegend | 304002 | Barcode |
| 106Pd | CD45 | HI30 | Biolegend | 304002 | Barcode |
| 113In | CD61 | VI-PL2 | Biolegend | 336402 | Pregate |
| 115In | CD11c | Bu15 | Biolegend | 337202 | Phenotype |
| 139La | FOXP3 | 22510 | Abcam | AB22510 | Activity |
| 141Pr | CD20 | 2H7 | Biolegend | 302302 | Phenotype |
| 142Nd | CD19 | HIB19 | Biolegend | 302202 | Phenotype |
| 143Nd | CD45RA | HI100 | Biolegend | 304102 | Phenotype |
| 144Nd | CD80 | EPR1157(2) | Abcam | AB224797 | Activity |
| 145Nd | CD8a | RPA-T8 | Biolegend | 301002 | Phenotype |
| 146Nd | CD24 | ML5 | Biolegend | 311103 | Phenotype |
| 147Sm | CXCR5 (CD185) | RF8B2 | BD | 552032 | Activity |
| 148Nd | CD5 | UCHT2 | Biolegend | 300602 | Phenotype |
| 149Sm | CD45RO | UCHL1 | Biolegend | 304239 | Phenotype |
| 150Nd | CD138 | DL-101 | Biolegend | 352302 | Phenotype |
| 151Eu | CD22 | HIB22 | Biolegend | 302502 | Activity |
| 152Sm | CD21 | BU32 | Biolegend | 354902 | Phenotype |
| 153Eu | CD66b | G10F5 | BD | 555723 | Pregate |
| 154Sm | p38 MAPK | 30/p38 MAPK | BD | 612281 | Activity |
| 155Gd | CD27 | M-T271 | BD | 555439 | Phenotype |
| 156Gd | CD10 | HI10a | Biolegend | 312202 | Phenotype |
| 158Gd | CD40 | 5C3 | Biolegend | 334325 | Activity |
| 159Tb | CD235ab | HIR2 | Biolegend | 306602 | Pregate |
| 160Gd | Igκ | JDC-12 | BD | 555789 | Activity |
| 161Dy | CD62L | DREG-56 | Biolegend | 304835 | Phenotype |
| 163Dy | CXCR3 (CD183) | REA232 | Miltenyi | 130-095-212 | Activity |
| 165Ho | IgG | G18-145 | BD | 555784 | Phenotype |
| 166Er | p65 NFκB | K10-895.12.50 | BD | 558393 | Activity |
| 167Er | CD38 | HIT2 | Biolegend | 303502 | Phenotype |
| 168Er | CD4 | SK3 | Biolegend | 344602 | Phenotype |
| 169Tm | CD25 | M-A251 | Biolegend | 356102 | Phenotype |
| 170Er | CD3 | UCHT1 | Biolegend | 300402 | Phenotype |
| 171Yb | Blimp-1 | ROS195G | Biolegend | 648202 | Activity |
| 172Yb | IgM | G20127 | BD | 555780 | Phenotype |
| 173Yb | MAPKAPK2 | P24-694 | BD | 562469 | Activity |
| 174Yb | G6 [IGHV1-69][A] | G6 | n/a | n/a | Activity |
| 175Lu | PD-1 (CD279) | EH12.2H7 | Biolegend | 329902 | Activity |
| 176Yb | CD127 | A019D5 | Biolegend | 351302 | Phenotype |

All antibodies were metal-conjugated in-house at Sydney Cytometry (Sydney, NSW). [A]G6 antibody was provided by A/Prof Joanne Reed, Westmead Institute for Medical Research (Sydney, NSW).

**Supplementary Table 2.** CyTOF antibodies used for further assessment of circulating B cell subpopulations during the follow-up mass cytometry study.

| Label | Target | Clone | Supplier | Catalogue # | Use |
|---|---|---|---|---|---|
| 89Y | CD8a | RPA-T8 | Biolegend | 301002 | Phenotype |
| 104Pd | CD45 | HI30 | Biolegend | 304002 | Barcode |
| 106Pd | CD45 | HI30 | Biolegend | 304002 | Barcode |
| 113In | CD61 | VI-PL2 | Biolegend | 336402 | Pregate |
| 115In | Ki67 | B56 | BD | 556003 | Activity |
| 141Pr | CD20 | 2H7 | BD | 555677 | Phenotype |
| 142Nd | CD19 | HIB19 | Biolegend | 302202 | Phenotype |
| 143Nd | CD5 | UCHT2 | Biolegend | 300602 | Phenotype |
| 144Nd | CD86 | IT2.2 | Biolegend | 305402 | Activity |
| 145Nd | CD4 | RPA-T4 | BD | 555344 | Phenotype |
| 147Sm | CD95L (FasL) | NOK-1 | Biolegend | 306402 | Activity |
| 150Nd | CD138 | DL-101 | Biolegend | 352302 | Phenotype |
| 151Eu | CD22 | HIB22 | Biolegend | 302511 | Activity |
| 152Sm | CD21 | Bu32 | Biolegend | 354902 | Phenotype |
| 153Eu | CD66b | G10F5 | BD | 555723 | Pregate |
| 154Sm | CD10 | HI10a | Biolegend | 312202 | Phenotype |
| 155Gd | CD27 | M-T271 | BD | 555439 | Phenotype |
| 156Gd | 9G4 [IGHV4-34][A] | 9G4 | n/a | n/a | Activity |
| 158Gd | FCRL4 (CD307d) | 413D12 | Biolegend | 340202 | Phenotype |
| 159Tb | CD235ab | HIR2 | Biolegend | 305402 | Pregate |
| 161Dy | PD-L1 (CD274) | 29E.2A3 | Biolegend | 329702 | Activity |
| 162Dy | CD80 | L307.4 | BD | 557223 | Activity |
| 165Ho | CD40 | 5C3 | Biolegend | 334302 | Activity |
| 166Er | IgD | IA6-2 | BD | 555776 | Phenotype |
| 167Er | CD38 | HIT2 | Biolegend | 303502 | Phenotype |
| 168Er | CD95 (FasR) | DX2 | Biolegend | 305602 | Activity |
| 169Tm | FOXP3 | PCH101 | eBioscience | 14-4776-82 | Activity |
| 170Er | CD3 | UCHT1 | Biolegend | 300402 | Phenotype |
| 171Yb | CD23 (FcεRII) | EBVCS-5 | Biolegend | 338502 | Activity |
| 174Yb | G6 [IGHV1-69][A] | G6 | n/a | n/a | Activity |
| 175Lu | PD-1 (CD279) | EH12.2H7 | Biolegend | 329902 | Activity |
| 176Yb | IgM | MHM-88 | Biolegend | 314502 | Phenotype |

All antibodies were metal-conjugated in-house at Sydney Cytometry (Sydney, NSW). [A]The G6 and 9G4 antibodies were provided by A/Prof Joanne Reed, Westmead Institute for Medical Research (Sydney, NSW).

**Supplementary Table 3.** Summary of steps in the mass cytometry staining protocol.

| Step | Description | Buffer used |
|---|---|---|
| *Day 1: Lysis and surface staining* | | |
| Thawing | Remove first batch of samples from -80°C storage and transfer to 4°C to thaw for approximately 45 mins. | n/a |
| Red blood cell lysis | Lyse cells in 1:1000 Thaw-Lyse Buffer 1 (Smart Tube Inc.), then in 1:5 Lyse Buffer 2 (Smart Tube Inc.) according to manufacturer's instructions. | Diluted in MilliQ®-purified water. |
| Cell counting | Count cells with haemocytometer, then transfer $2x10^6$ cells per sample for staining. Transfer excess to a new tube to use as a surface gating control. | FACS |
| Heparin block | Block in 200 µL 100IU/mL Heparin for 20 mins at RT. | FACS |
| Barcoding | Incubate with unique CD45 barcode solution for 30 mins at 4°C, wash twice in FACS, then combine barcoded pairs. | FACS |
| Fc receptor block | Block in 5 µL human Fc-receptor blocking solution (Cell Signalling Technology) for 10 mins at RT. | 0.5% FBS in 1X PBS |
| Surface staining[A] | Incubate with surface antibodies for 30 mins at 4°C. | FACS |
| Fixation/permeabilization | Resuspend in 1:3 Fix/Perm buffer for 30 mins at RT, then leave at 4°C overnight. | Fix/Perm |
| *Day 2: Intracellular and nuclear staining* | | |
| Washing | Wash twice in FACS, then once in Perm. | FACS, Perm |
| Heparin block | Block in 200 µL 100IU/mL Heparin for 20 mins at RT. | Perm |
| Intracellular staining | Incubate with intracellular antibodies for 30 mins at 4°C. | Perm |
| Methanol permeabilization | Incubate with 500 µL ice-cold 100% methanol (added dropwise while vortexing tube) for 30 mins at 4°C. | Methanol |
| Heparin block | Block in 200 µL 100IU/mL Heparin for 20 mins at RT. | FACS |
| Nuclear antibody staining | Incubate with nuclear antibodies for 30 mins at 4°C. | FACS |
| DNA intercalator | Incubate in 1:4000 DNA-Ir in 4% paraformaldehyde for 20 mins at RT, then 4°C overnight. | PBS |
| *Day 3-5: Acquisition* | | |
| Washing | Wash once in FACS, then once in MilliQ®-purified water. | FACS |
| Cell counting | Resuspend cells in CAS and count with haemocytometer. Centrifuge, aspirate, keep pallet at 4°C until acquisition. | CAS |
| Resuspension | Resuspend at $1x10^6$ cells/mL in CAS containing 11% EQ beads, then filter twice through a cell strainer cap. | CAS |
| Acquisition | Run samples on Helios mass cytometer. Normalise signal intensities to EQ beads after acquisition. | n/a |

After each antibody incubation, Heparin blocking step, and methanol permeabilization, cells were topped up with 1mL of the relevant staining buffer and centrifuged at 4˚C (600 g for 5 mins before fixation/permeabilization, or 800 g for 7 mins after fixation/permeabilization). [A]Surface Ig markers were added sequentially to reduce cross-reactivity. CAS = Maxpar® Cell Acquisition Solution (Standard BioTools); DNA-Ir = Cell-ID™ DNA Intercalator-Ir (Standard BioTools); EQ = EQ™ Four Element Calibration Beads (Standard BioTools); FACS = eBioscence™ Flow Cytometry Staining Buffer (Invitrogen); Fix/Perm = eBioscience™ Fixation/Permeabilization Buffer (Invitrogen); Perm = eBiosience™ Permeabilization Buffer (Invitrogen).

**Supplementary Table 4.** Classification of major circulating B cell and T cell subpopulations based on core surface marker expression.

| CD19+ B cells | IgD | IgM | CD27 | CD21 | CD38 | CD20 | CD10 | CD138 |
|---|---|---|---|---|---|---|---|---|
| Transitional 1 | +/lo | hi | – | lo | hi | hi | hi | – |
| Transitional 2 | + | + | – | + | + | ++ | + | – |
| Transitional 3 | + | + | – | + | + | + | lo/– | – |
| Naive[A] | + | + | – | + | int | + | – | – |
| Unswitched memory[A] | + | hi | int | + | lo | + | – | – |
| Marginal zone | lo/– | + | int | + | lo | + | – | – |
| Class-switched memory[A] | – | – | + | + | – | + | – | – |
| DN1/2 | – | – | – | +/– | +/– | + | – | – |
| Unswitched plasmablast | +/– | +/– | lo/– | lo/– | ++ | lo | – | – |
| Class-switched plasmablast | – | – | + | – | hi | lo/– | – | lo |
| Plasma cell | – | – | ++ | – | hi | – | – | hi |
| **CD3+ T cells** | **CD4** | **CD8a** | **CD25** | **FOXP3** | **CD62L** | **CD45RA** | **CD45RO** | **CD27** |
| Helper | + | – | – | – | | | | |
| Cytotoxic | – | + | – | – | | | | |
| Regulatory | + | – | + | + | | | | |
| Naive | | | | | hi | + | – | + |
| Effector | | | | | – | + | – | – |
| Effector memory | | | | | – | – | + | +/– |
| Central memory | | | | | + | – | + | hi |

[A]Activated subsets defined as $CD21^{-}$. DN = double-negative.

**Supplementary Table 5.** FlowSOM clustering parameters for unsupervised CyTOF analysis.

| Pregate | Clusters | Clusters used | Merged metaclusters | Markers used for clustering and dimensionality reduction |
|---|---|---|---|---|
| $CD19^{+}$ | 10x10=100 | 24/30 | 21 | CD20, IgD, IgM, CD38, CD27, CD21, CD138, CD95, CD86, CD3 |
| $CD3^{+}$ | 10x10=100 | 16/25 | 16 | CD4, CD8a, CD25, CD27, CD62L, CD45RA, CD45RO, CD19 |

**Supplementary Table 6.** Final CRPS and control cohort demographics across all studies.

| | Female/male | | | Age (years) | | |
|---|---|---|---|---|---|---|
| | *CRPS* | *Control* | *P-value*[A] | *CRPS* | *Control* | *P-value*[B] |
| Preliminary CyTOF | 20/6 | 18/6 | >0.9999 | 45.2 ± 14.4 | 43.7 ± 13.7 | 0.7508 |
| Follow-up CyTOF | 31/10 | 30/10 | >0.9999 | 45.7 ± 14.8 | 42.9 ± 13.2 | 0.3806 |
| Serum proteomics | 31/11 | 15/5 | >0.9999 | 45.8 ± 14.7 | 45.6 ± 11.8 | 0.9527 |
| Autoantibody screen | 31/11 | 13/5 | >0.9999 | 45.8 ± 13.7 | 45.6 ± 12.1 | 0.9679 |

Data are *n* values or mean ± SD as appropriate. [A]Fisher's exact tests; [B]Welch's t-tests.

**Supplementary Table 7.** CRPS participant demographics and clinical information.

| Subject | Sex | Age | Duration | Inciting injury | CRPS-affected area | Current medications[A] | Comorbidites |
|---|---|---|---|---|---|---|---|
| 1 | F | 37 | 14 years | Left knee dislocation | Left lower limb, right lower, left upper limb | Baclofen, duloxetine, pregabalin | None |
| 2 | M | 48 | 7 months | Left radius/ulnar fracture | Left upper limb | Amitriptyline, gabapentin | None |
| 3 | M | 55 | 7 years | Three surgeries in left wrist after ligament tear | Left upper limb, possible right upper | Clonidine, dosulepin, gabapentin, ketamine infusion | Left ankle pain from bone coalition |
| 4 | F | 50 | 3.5 years | Brachial plexus injury | Right upper limb, possible right lower | Buprenorphine, nortriptyline, topiramate | None |
| 5 | M | 34 | 14 years | Right hand amputation after car accident | Right upper limb, left upper limb, both lower limbs | Fluorometholone, gabapentin, medicinal cannabis, nortriptyline, sodium valproate, zopiclone | Phantom limb pain; neck/head pain |
| 6 | M | 20 | 1.5 years | Right foot crush injury with fracture | Right lower limb | Pregabalin, venlafaxine | None |
| 7 | M | 32 | 13 years | Right shoulder injury | Right upper limb, possible left upper | Buprenorphine patch, clonidine, domperidone, esomeprazole, gabapentin, prednisolone, quetiapine, sodium valproate, Symbicort (budesonide/formoterol), tapentadol, Ventolin (salbutamol), zopiclone | Neurofibromatosis, bipolar disorder, PTSD |
| 8 | F | 33 | 27 months | Right ankle fracture and surgery | Right lower limb, possible left lower | Diazepam, gabapentin, oxycodone propanolol, phentermine, tapentadol | None |
| 9 | F | 78 | 3 years | Left hand surgery | Left upper limb (hand) | Paracetamol/codeine | Osteoporosis |
| 10 | F | 42 | 7 years | Right ankle fracture | Right lower limb, possible right upper | Escitalopram, esomeprazole, Flixotide (fluticasone proprionate), paracetamol, Ventolin (salbutamol) | Fainting syndrome |
| 11 | F | 66 | 12 years | Shoulder surgery | Left upper limb (shoulder/elbow/hand) | Atenolol, duloxetine, famciclovir, meloxicam, paracetamol/codeine | COPD, fibromyalgia, long COVID |
| 12 | F | 60 | 6 years | Comminuted fracture in left ankle | Left lower limb (ankle) | None | None |
| 13 | F | 26 | 13 years | Right elbow injury, ulnar nerve release, shoulder injury/reconstruction. | Right upper limb (shoulder/arm), right lower-mid back | Gabapentin, midrodine | POTS |
| 14 | F | 44 | 8.8 years | Three surgeries after left foot Lisfranc fracture | Left lower limb | Anti-depressant, ketamine infusion, paracetamol, pregabalin, tramadol | Lyme disease |
| 15 | F | 59 | 5 months | Dupuytren's hand surgery | Right upper limb (hand/arm/shoulder) | Mersyndol (paracetamol/codeine/ doxylamine succinate) | None |
| 16 | F | 38 | 8 years | Right shoulder subluxation | Right upper limb, right lower limb | Clonidine, drospirenone, duloxetine, melatonin, nizatidine, topiramate | Endometriosis, fibromyalgia, GERD |
| 17 | F | 46 | 14 months | Right elbow/knee fractures | Right lower limb, left lower limb | Paracetamol/codeine, quetiapine, rosuvastatin, tapentadol, venlafaxine | None |
| 18 | M | 63 | 3 years | Right knee replacement and corrective surgery | Right lower limb (knee), lower back | Duloxetine, gabapentin | None |

| | | | | | | | |
|---|---|---|---|---|---|---|---|
| 19 | F | 27 | 1 year | Left knee dislocation | Left lower limb | Pregabalin, tapentadol | None |
| 20 | F | 59 | 9 months | Radial fracture in left arm | Right upper limb (wrist/arm/shoulder), right knee | Naproxen, pregabalin | None |
| 21 | F | 49 | 6 years | Left foot fracture and ligament tear with reconstructive surgery | Left lower limb (foot) | Clonidine, duloxetine, esomeprazole, ibuprofen, naltrexone, pregabalin, topiramate | Endometriosis |
| 22 | F | 22 | 3 years | Right hand tendon injury | Right upper limb | Celecoxib, lidocaine patch, tapentadol | None |
| 23 | M | 43 | 3 years | Left foot fracture | Left lower limb (foot) | Not collected | Not collected |
| 24 | F | 42 | 12 years | Two surgeries after right foot crush injury with Lisfranc fracture | Right lower limb (foot), widespread pain | Melatonin, oxycodone, promethazine | None |
| 25 | F | 51 | 2 years | Left distal radius fracture | Left upper limb (hand/wrist) right hand, left/right foot | Amitriptyline, gabapentin | None |
| 26 | F | 37 | 10 years | Partial dislocation of left shoulder | Left upper limb, possible left lower limb | Chlorpromazine, diazepam, fluoxetine, methadone, oxycodone. | None |
| 27 | F | 62 | 3 months | Right wrist fracture/ sprain | Right upper limb (wrist) | Gabapentin, prednisolone, tapentadol | None |
| 28 | F | 19 | 27 months | Soft tissue injury | Lower limb | Not collected | Not collected |
| 29 | M | 22 | 25 months | Fracture | Lower limb | Not collected | Not collected |
| 30 | F | 48 | 2.5 years | Fracture | Lower limb | Not collected | Not collected |
| 31 | M | 37 | 5 months | Fracture | Lower limb | Not collected | Not collected |
| 32 | M | 32 | 3 months | Fracture | Lower limb | Not collected | Not collected |
| 33 | F | 58 | 3.5 years | Surgery | Lower limb | Not collected | Not collected |
| 34 | M | 53 | 5.5 years | Fracture | Lower limb | Not collected | Not collected |
| 35 | F | 44 | 4 years | Fracture | Upper limb | Not collected | Not collected |
| 36 | F | 59 | 3.2 years | Surgery | Upper limb | Not collected | Not collected |
| 37 | F | 61 | 24.4 years | Soft tissue injury | Upper limb | Not collected | Not collected |
| 38 | F | 57 | 3.3 years | Fracture | Upper limb | Not collected | Not collected |
| 39 | F | 53 | 6 months | Soft tissue injury | Upper limb | Not collected | Not collected |
| 40 | F | 25 | 6 months | Soft tissue injury | Upper limb | Not collected | Not collected |
| 41 | F | 61 | 5.3 years | Soft tissue injury | Upper limb | Not collected | Not collected |
| 42 | F | 70 | 4 months | Surgery | Upper limb | Not collected | Not collected |

Table shows sex, age, duration (time since onset), inciting injury, CRPS-affected region (primary area underlined), current medications, and comorbidities reported by each CRPS participant. Samples from subjects 1-27 were collected between 2022 and 2024 at the Baker Heart and Diabetes Institute (Melbourne, NSW) and were first profiled in the preliminary CyTOF study then in the follow-up CyTOF study, proteomic screen, and autoantibody assay, in addition to samples from subjects 28-42, which were collected between 2017 and 2018 at Genesis Research Services (Broadmeadow, NSW). [A]Participants were asked to cease any immunomodulating medications (NSAIDs, opioids, and corticosteroids) for 7 days prior to blood collection ($n$ = 39 complied). COPD = chronic obstructive pulmonary disease; FMS = fibromyalgia syndrome; GERD = gastroesophageal reflux disease; POTS = postural orthostatic tachycardia syndrome; PTSD = post-traumatic stress disorder.

**Supplementary Table 8.** Full surface phenotypes for the FlowSOM metaclusters identified by unsupervised analysis of pre-gated CD19$^{+}$ and CD3$^{+}$ compartments.

| **Name** | **Phenotype** |
|---|---|
| *CD19$^{+}$ B cell metaclusters (all CD3$^{-}$)* | |
| Transitional 1 | IgM$^{+}$IgD$^{-}$CD27$^{-}$CD21$^{-}$CD38$^{+}$CD95$^{lo}$CD86$^{lo}$CD20$^{+}$ |
| Transitional 2/3 | IgM$^{+}$IgD$^{+}$ CD27$^{-}$CD21$^{+}$CD38$^{++}$CD95$^{+}$CD86$^{-}$CD20$^{+}$ |
| Naive | IgM$^{+}$IgD$^{+}$CD27$^{-}$CD21$^{+}$CD38$^{int}$CD95$^{-}$CD86$^{-}$CD20$^{+}$ |
| Naive activated | IgM$^{+}$IgD$^{+}$CD27$^{-}$CD21$^{-}$CD38$^{lo}$CD95$^{-}$CD86$^{-}$CD20$^{+}$ |
| Naive activated IgMIgDlo | IgM$^{lo}$IgD$^{lo}$CD27$^{-}$CD21$^{-}$CD38$^{+}$CD95$^{+}$CD86$^{-}$CD20$^{+}$ |
| Naive IgMlo | IgM$^{-}$IgD$^{+}$CD27$^{-}$CD21$^{+}$CD38$^{int}$CD95$^{-}$CD86$^{-}$CD20$^{int}$ |
| Unswitched memory | IgM$^{+}$IgD$^{+}$CD27$^{lo}$CD21$^{+}$CD38$^{int}$CD95$^{lo}$CD86$^{+}$CD20$^{+}$ |
| Marginal zone CD21lo | IgM$^{+}$IgD$^{lo}$CD27$^{+}$CD21$^{-}$CD38$^{-}$CD95$^{lo}$CD86$^{+}$CD20$^{+}$ |
| CSM resting | IgM$^{-}$IgD$^{-}$CD27$^{+}$CD21$^{+}$CD38$^{-}$CD95$^{-}$CD86$^{-}$CD20$^{+}$ |
| CSM intermediate | IgM$^{-}$IgD$^{-}$CD27$^{+}$CD21$^{+}$CD38$^{int}$CD95$^{+}$CD86$^{+}$CD20$^{+}$ |
| CSM activated | IgM$^{-}$IgD$^{-}$CD27$^{+}$CD21$^{-}$CD38$^{-}$CD95$^{+}$CD86$^{+}$CD20$^{+}$ |
| DN1 | IgM$^{-}$IgD$^{-}$CD27$^{-}$CD21$^{+}$CD38$^{lo}$CD95$^{-}$CD86$^{-}$CD20$^{+}$ |
| DN2-like CD38lo | IgM$^{-}$IgD$^{-}$CD27$^{-}$CD21$^{-}$CD38$^{lo}$CD95$^{-}$CD86$^{-}$CD20$^{+}$ |
| DN2 | IgM$^{lo}$IgD$^{-}$CD27$^{-}$CD21$^{-}$CD38$^{-}$CD95$^{-}$CD86$^{-}$CD20$^{+}$ |
| Plasmablast | IgM$^{-}$IgD$^{-}$CD27$^{+}$CD21$^{-}$CD38$^{hi}$CD95$^{++}$CD86$^{-}$CD20$^{lo}$CD138$^{lo}$ |
| Plasmablast activated | IgM$^{int}$IgD$^{-}$CD27$^{+}$CD21$^{-}$CD38$^{hi}$CD95$^{hi}$CD86$^{++}$CD20$^{+}$CD138$^{+}$ |
| Naive CD20lo | IgM$^{+}$IgD$^{+}$CD27$^{-}$CD21$^{+}$CD38$^{lo}$CD95$^{-}$CD86$^{-}$CD20$^{lo}$ |
| Naive activated CD20lo | IgM$^{+}$IgD$^{+}$ CD27$^{-}$CD21$^{-}$CD38$^{-}$CD95$^{-}$CD86$^{-}$ CD20$^{lo}$ |
| Marginal zone CD20lo | IgM$^{+}$IgD$^{-}$CD27$^{lo}$CD21$^{+}$CD38$^{-}$CD95$^{-}$CD86$^{-}$CD20$^{lo}$ |
| CSM resting CD20lo | IgM$^{lo}$IgD$^{-}$CD27$^{+}$CD21$^{+}$CD38$^{-}$CD95$^{-}$CD86$^{-}$CD20$^{lo}$ |
| DN CD20lo | IgM$^{-}$IgD$^{-}$CD27$^{-}$CD21$^{-}$CD38$^{+}$CD95$^{+}$CD86$^{lo}$CD20$^{lo}$ |
| *CD3$^{+}$ T cell metaclusters (all CD19$^{-}$)* | |
| Th naive | CD4$^{+}$CD8a$^{-}$CD25$^{-}$CD62L$^{+}$CD45RA$^{+}$CD45RO$^{-}$CD27$^{+}$ |
| Th effector | CD4$^{+}$CD8a$^{-}$CD25$^{-}$CD62L$^{lo}$CD45RA$^{+}$CD45RO$^{-}$CD27$^{-}$ |
| Th intermediate | CD4$^{+}$CD8a$^{-}$CD25$^{-}$CD62L$^{lo}$CD45RA$^{-}$CD45RO$^{-}$CD27$^{-}$ |
| Th effector memory | CD4$^{+}$CD8a$^{-}$CD25$^{-}$CD62L$^{lo/-}$CD45RA$^{-}$CD45RO$^{+}$CD27$^{-}$ |
| Th Temra | CD4$^{+}$CD8a$^{-}$CD25$^{-}$CD62L$^{lo}$CD45RA$^{+}$CD45RO$^{+}$CD27$^{lo}$ |
| Th central memory | CD4$^{+}$CD8a$^{-}$CD25$^{-}$CD62L$^{lo}$CD45RA$^{-}$CD45RO$^{+}$CD27$^{+}$ |
| Treg naive | CD4$^{+}$CD8a$^{-}$CD25$^{+}$CD62L$^{+}$CD45RA$^{+}$CD45RO$^{-}$CD27$^{+}$ |
| Treg effector memory | CD4$^{+}$CD8a$^{-}$CD25$^{+}$CD62L$^{-}$CD45RA$^{-}$CD45RO$^{+}$CD27$^{lo}$ |
| Tc naive | CD4$^{-}$CD8a$^{+}$CD25$^{-}$CD62L$^{+}$CD45RA$^{+}$CD45RO$^{-}$CD27$^{+}$ |
| Tc effector | CD4$^{-}$CD8a$^{+}$CD25$^{-}$CD62L$^{lo}$CD45RA$^{+}$CD45RO$^{-}$CD27$^{-}$ |
| Tc effector memory | CD4$^{-}$CD8a$^{+}$CD25$^{-}$CD62L$^{+}$CD45RA$^{-}$CD45RO$^{+}$CD27$^{+}$ |
| Tc Temra | CD4$^{-}$CD8a$^{+}$CD25$^{-}$CD62L$^{+}$CD45RA$^{lo}$CD45RO$^{lo}$CD27$^{+}$ |
| CD4+CD8a+ naive | CD4$^{+}$CD8a$^{+}$CD25$^{-}$CD62L$^{+}$CD45RA$^{+}$CD45RO$^{-}$CD27$^{+}$ |
| CD4+CD8a+ memory | CD4$^{+}$CD8a$^{+}$CD25$^{-}$CD62L$^{lo}$CD45RA$^{-}$CD45RO$^{+}$CD27$^{-}$ |
| CD4-CD8a- naive | CD4$^{-}$CD8a$^{-}$CD25$^{-}$CD62L$^{lo}$CD45RA$^{+}$CD45RO$^{-}$CD27$^{-}$ |
| CD4-CD8a- memory | CD4$^{-}$CD8a$^{-}$CD25$^{-}$CD62L$^{-}$CD45RA$^{-}$CD45RO$^{+}$CD27$^{-}$ |

**Supplementary Methods 1.** CyTOF staining and acquisition.

*Lysis and staining.* On the first day of staining for each batch, whole blood samples were removed from -80˚C storage and transferred to 4˚C to thaw for approximately 45 mins. Erythrocyte lysis was performed using Thaw Lyse buffer 1 (1:1000 in MilliQ®-purified water; SmartTube Inc.) and Lyse buffer 2 (1:5 in MilliQ®-purified water; SmartTube Inc.) according to the manufacturer's instructions, until the cell pellet was pink-white. Cells were counted using a haemocytometer, and $2x10^6$ per sample transferred for staining. Excess cells were added to a new tube for a gating control (surface stain only).

All antibodies were metal-conjugated in-house at Sydney Cytometry (Sydney, NSW). Antibody cocktails were prepared using eBioscience™ FACS buffer for surface and nuclear antibodies, or eBioscience™ Perm buffer (1:10 in MilliQ®-purified water) for intracellular antibodies. All antibody incubations were performed at 100 µL per $2x10^6$ cells for 30 mins at 4˚C. Before each staining step, cells were incubated in 100IU/mL Heparin in the relevant staining buffer for 20 mins at RT to block non-specific eosinophil staining [5]. After each Heparin blocking step or antibody incubation, cells were washed in the relevant staining buffer and centrifuged at 4˚C at 600 g for 5 mins (before fixation/permeabilization) or 800 g for 7 mins (after fixation/permeabilization).

Isolated PBMCs were first incubated in a unique CD45 barcode solution (104Pd-CD45 for CRPS; 106Pd-CD45 for controls), allowing samples to be combined such that each new tube contained cells from one CRPS participant and one matched control subject. Cells were washed twice before combining samples (to minimise barcode crossover). After barcoding, cells were incubated with 5 µL human Fc-receptor blocking solution (in PBS + 0.5% FBS; Cell Signalling Technology) for 10 mins at RT, then stained with surface antibodies. Surface Ig markers were separated into two sequential staining steps to reduce cross-reactivity. Cells were then incubated in eBioscience™ Fix/Perm buffer (1:3 concentrate:diluent) for 30 mins at RT, then kept at 4˚C overnight. The next day, cells were washed twice in FACS, once in Perm, then stained with intracellular antibodies. To prepare for nuclear staining, cells were incubated in 500 µL ice-cold 100% methanol (added dropwise while vortexing tube) for 30 mins at 4˚C, then washed in FACS. Cells were stained with nuclear antibodies, then fixed in 4% paraformaldehyde in PBS containing 1:4000 Cell-ID™ DNA-Intercalator-Ir (Standard BioTools) for 20 mins at RT, then kept at 4˚C until acquisition.

*Sample acquisition.* On the day of acquisition, cells were washed once in FACS, once in MilliQ®-purified water, then resuspended in 1 mL Maxpar® Cell Acquisition Solution (CAS; Standard BioTools) and counted using a haemocytometer. Samples were then centrifuged at 800 g for 7 mins at 4˚C, aspirated, and kept at 4˚C until acquisition. Immediately before acquisition, pellets were resuspended at $1x10^6$ cells/mL in CAS containing 11% EQ™ Four Element calibration beads (Standard BioTools) and filtered twice through a cell strainer cap. All samples were acquired on a Helios suspension mass cytometer (Fluidigm). Signal intensities were normalised to EQ beads at 100 intervals/sec.

**Supplementary Methods 2.** Proteomic SWATH-MS acquisition and data analysis.

*Sample preparation.* Serum samples were processed as previously described [6]. In brief, total serum protein was quantified with the Pierce BCA Protein Assay Kit (Thermo Fisher Scientific, MA, USA). Equal amounts of protein from each sample were reduced with 15 mM dithiothreitol at 60˚C for 30 min, then alkylated with 30 mM iodoacetamide for 30 min at room temperature in the dark. The samples were subsequently digested overnight (16h) at 37˚C with trypsin at an enzyme-to-protein ratio of 1:30 (w/w) under gentle agitation. The resulting peptides were desalted and purified on C18 StageTips [7] prior to LC-MS/MS analysis.

*Protein analysis by LC-MS.* The desalted samples were reconstituted with 0.1% formic acid and analysed by liquid chromatography-mass spectrometry (LC-MS) as previously described [6]. Separation was performed on a Nexera series LC-40 UHPLC system (Shimadzu, Kyoto, Japan) fitted with an InfinityLab Poroshell 120 EC-C18 column (2.1 × 50 mm, 1.9 µm) and a matched UHPLC guard cartridge (Agilent, Santa Clara, CA, USA), maintained at 40 °C and coupled to a ZenoTOF 7600 (Sciex, Toronto, Canada). Mobile phase A was 0.1% (v/v) formic acid in water and mobile phase B was 0.1% (v/v) formic acid in acetonitrile. Peptides were eluted at 0.8 mL/min using a linear gradient from 9% to 29% B over 5.6 min, 29% to 42% B over 0.6 min, and 42% to 80% B over 0.2 min. The column was then washed at 80% B for 0.2 min and re-equilibrated at 3% B. Data was acquired in positive ion mode using sequential window acquisition of all theoretical (SWATH). Ion source was set to 1:60 psi, ion curtain gas to 40 psi and CAD gas to 7 psi. The ion spray voltage was 4500 V and the source temperature was 700°C. Ions were generated by electrospray and fragment ion spectra acquired across 60 overlapping windows spanning a precursor mass range of 400–900 m/z. Window widths ranged from approximately 5 to 42 m/z inclusive of a 1 m/z overlap and narrowed toward 650 m/z. The effective isolation windows were being 399.5-441.2, 440.2-459.9, etc. SWATH MS2 spectra were collected from 100 to 1800 m/z at an accumulation time of 5 ms per window in high-sensitivity mode with Zeno trap pulsing enabled, giving a total duty cycle of ~0.627 s. Collision energy for each window was ramped from 19 V to 43 V, at 1-2 V per window with increasing m/z.

*Proteomic data analysis.* Raw DIA-MS files were analysed using Spectronaut v20 (Biognosys) with the library-free directDIA workflow. Trypsin was specified as the digestion enzyme, allowing a maximum of one missed cleavage and a peptide length of 9-52 amino acids. Carbamidomethylation of cysteine was set as a fixed modification, while methionine oxidation and protein N-terminal acetylation were set as variable modifications, with up to five modifications permitted per peptide. Protein quantification required at least one unique peptide per protein. The false discovery rate (FDR) was controlled at 1% at both the peptide and protein levels. Cross-run normalisation was performed within Spectronaut. Protein groups were retained for analysis if they were quantified in at least 50% of samples across the combined cohort [8,9]. Missing values, which in mass spectrometry-based proteomics commonly reflect protein abundances below the limit of detection, were imputed under a missing-not-at-random assumption [10]. Relative protein abundances were compared between the cohorts using a t-test in RStudio. Proteins with a P-value below 0.05 and a fold change greater than 1.5 were considered differentially expressed.

### Supplementary material references


1 LOVIBOND, S. H. & LOVIBOND, P. F. 1995. *Manual for the Depression Anxiety Stress Scales,* 2nd Ed. Sydney, Australia: Psychology Foundation of Australia.

2 SULLIVAN, M. J. L. 1995-2009. T*he Pain Catastrophizing Scale - User Manual.* Montreal, Quebec: McGill University.

3 NICHOLAS, M. K. 2007. The pain self-efficacy questionnaire: Taking pain into account. *Eur J Pain,* 11**,** 153-63.

4 NEBLETT, R., HARTZELL, M. M., MAYER, T. G., BRADFORD, E. M. & GATCHEL, R. J. 2016. Establishing clinically meaningful severity levels for the Tampa Scale for Kinesiophobia (TSK-13). *Eur J Pain,* 20**,** 701-10.

5 RAHMAN, A.H., TORDESILLAS, L. & BERIN, M.C. 2016. Heparin reduces nonspecific eosinophil staining artifacts in mass cytometry experiments. *Cytometry A,* 89**,** 601-7.

6 HENG, B., GUNASEGARAN, B., KRISHNAMURTHY, S., BUSTAMANTE, S., PIRES, A.S., CHOW, S., AHN, S.B., PAUL-HENG, M., MACIVER, Y., SMITH, K., TRAN, D.P., HOWLEY, P.P., BILGIN, A.A., SHARLAND, A., SCHLOEFFEL, R. & GUILLEMIN, G.J. 2025. Mapping the complexity of ME/CFS: Evidence for abnormal

energy metabolism, altered immune profile, and vascular dysfunction. *Cell Rep Med,* 6, 102514.

7 RAPPSILBER, J., MANN, M. & ISHIHAMA, Y. 2007. Protocol for micro-purification, enrichment, pre-fractionation and storage of peptides for proteomics using StageTips. *Nat Protoc,* 2**,** 1896-906.

8 LAZAR, C., GATTO, L., FERRO, M., BRULEY, C. & BURGER, T. 2016. Accounting for the Multiple Natures of Missing Values in Label-Free Quantitative Proteomics Data Sets to Compare Imputation Strategies. *J Proteome Res,* 15**,** 1116-25.

9 MCCLINTICK, J.N. & EDENBERG, H.J. 2006. Effects of filtering by Present call on analysis of microarray experiments. *BMC Bioinformatics,* 7**,** 49.

10 LIU, M. & DONGRE, A. 2021. Proper imputation of missing values in proteomics datasets for differential expression analysis. *Brief Bioinform,* 22.
